# DC-coupled, 2.5D electrophysiological imaging of large-scale cortical dynamics

**DOI:** 10.1101/2024.12.20.629545

**Authors:** R. Garcia-Cortadella, J. Cisneros-Fernandez, G. Schwesig, A. Shahidi, A. Umurzakova, N. Schäfer, J. Aguilar, E. Masvidal-Codina, E. Del Corro, R. Moahrlok, M. Kurnoth, J. Paetzold, H. Loeffler, C. Jeschke, J. Meents, X. Illa, F. Serra-Graells, A. Guimerà-Brunet, J.A. Garrido, A. Sirota

## Abstract

Infra-slow (<0.5 Hz) brain dynamics reflect homeostatic and neuromodulatory processes that modulate neuronal excitability and shape faster oscillations across brain regions. Infra-slow brain activity is typically inferred from magnetic or optical imaging, but these methods are limited in temporal resolution and compatibility with unconstrained behavior. Infra-slow local field potentials (isLFPs) could provide a direct measure of infra-slow network dynamics, but have remained poorly characterized due to the absence of scalable, DC-coupled recording methods. Here, we introduce DC-coupled electrophysiological imaging based on arrays of up to 512 multiplexed graphene transistors enabling stable, high-density recordings across cortical regions and cortical layers in freely moving rats. We developed an analytical framework for the analysis of wide-band LFP, revealing that synchronous oscillatory states of variable duration and spatial scale are consistently linked to topographically and translaminarly structured DC potential shifts. We propose a physiological model linking these DC shifts to sustained gradients of extracellular K^+^ concentration, providing a mechanistic connection between neuronal synchrony and isLFP dynamics. By integrating DC-coupled sensing, multiplexed scalability, and depth-surface co-registration, this work establishes a new modality for imaging-like electrophysiology in freely moving animals and a framework for interpreting infra-slow dynamics.

## Introduction

Large-scale neuronal dynamics provide the scaffold for neuronal communication and computation across spatiotemporal scales, coordinating the emergence of diverse behavioral and cognitive states^1–4^. Within this spatiotemporal hierarchy, infra-slow (<0.5 Hz) dynamic constitutes a fundamental component of functional brain activity, reflecting the interplay of neuronal, metabolic, and neuromodulatory processes^5–9^ that regulate cortical excitability and shape the emergence of faster oscillatory states. As such, infra-slow dynamics provide a temporal framework for cognitive processing, arousal and attentional states^10–15^ and its readout is invaluable for studying behavior and high-order functions.

Infra-slow activity has been inferred primarily from functional magnetic resonance imaging (fMRI)^3,16,17^ and optical imaging in head-fixed preparations^18–20^. Although these readout modalities reveal the large-scale spatial organization of infra-slow activity, they suffer from coarse temporal resolution and are incompatible with unconstrained behavior. Alternatively, DC-coupled electrophysiological recordings provide direct access to wide-band dynamics, revealing infra-slow local field potentials (isLFPs) that correlate with imaging signals^19,21,22^, while providing an insight into fast time scale brain dynamics. isLFP fluctuations are thought to arise from shifts in direct current (DC) potentials, which have been so far investigated in pathological brain states^23–30^. Large-scale mapping of isLFP patterns across physiological, behaviourally and functionally relevant brain states has been hindered by the lack of scalable, chronic DC-coupled recordings in freely-moving animals^31,32^. Consequently, the development of biophysical and analytical frameworks to understand the underlying mechanisms of their generation has also lagged behind^33^.

Resolving the physiological basis of infra-slow activity requires a method that combines large-scale topographical coverage, translaminar access, DC–to–high-frequency sensitivity, and stable long-term recordings in freely moving animals. Existing approaches, however, provide only subsets of these capabilities. Conventional DC-coupled systems rely on bulky electronics that do not scale to many channels and are highly susceptible to baseline drift and polarization artifacts^34,35^. Meanwhile, micro-electrocorticography (ECoG) arrays have been used to map mesoscale cortical dynamics, but their spatial extent and density are typically constrained by channel-count bottlenecks and surgical access^36–40^. Overcoming these technical challenges in freely-moving animals would open a unique window into anatomically-resolved behaviourally relevant infra-slow brain activity and its link to neuronal dynamics across temporal scales^4,6,12^, bridging insights from small animal models and non-invasive imaging in humans^10,21,22^.

To address these challenges, we developed a scalable DC-coupled electrophysiological recording platform based on large-scale multiplexed graphene solution-gated field-effect transistor (g-SGFETs) arrays. g-SGFETs provide electrochemically stable DC recording^30,41,42^ and support time-division multiplexing^43–45^, as demonstrated by previous implementations limited to small arrays and anesthetized or pathological conditions. Here, we demonstrate functional deployment of graphene bioelectronics for wide-band, naturalistic recordings through improvements in channel count, device uniformity, headstage miniaturization, extreme aspect-ratio grids and surgical scalability. Overcoming these limitations enabled chronic recordings with up to 512-channel multiplexed ECoG arrays and integration with multiplexed translaminar probes, resulting in 2.5-dimensional, DC-coupled electrophysiological imaging across cortical regions, layers, and behavioral states.

To gain a mechanistic insight into the generation and neuronal sources of isLFP, we developed an analytical framework to extract and relate the spatiotemporal structure of isLFP to faster brain dynamics. These recordings allowed precise identification of oscillatory states across spatial and temporal scales, revealing that synchronous thalamo-cortical oscillations are consistently associated with a specific topographic and translaminar DC potential pattern, contributing to the microstructure of sleep^46^. Furthermore, the generality and anatomical specificity of the observed DC pattern support a unified mechanistic model in which neuronal synchronization drives region- and layer-specific changes in extracellular K⁺ concentration, giving rise to the observed isLFP pattern. This model links slow and fast dynamics, reconciles prior imaging and electrophysiological findings, and provides mechanistic grounding for interpreting infra-slow activity. Together, these results establish 2.5D DC-coupled electrophysiological imaging as a scalable and mechanistically informative modality for investigating functional brain states across spatiotemporal scales.

## Results

### Scaling of multiplexed graphene probes and DC-coupled recording systems

Graphene transistors operate as transducers, converting extracellular potentials into variations in channel current (Fig. 1a), enabling high-quality DC-coupled recordings and implementation of time-domain multiplexing^44,45^. This architecture requires custom front-end electronics, including transimpedance amplifiers at the outputs of the array rows and a switching matrix to sequentially bias the source terminals of the transistors across the array columns (Fig. 1a).

**Figure 1.**
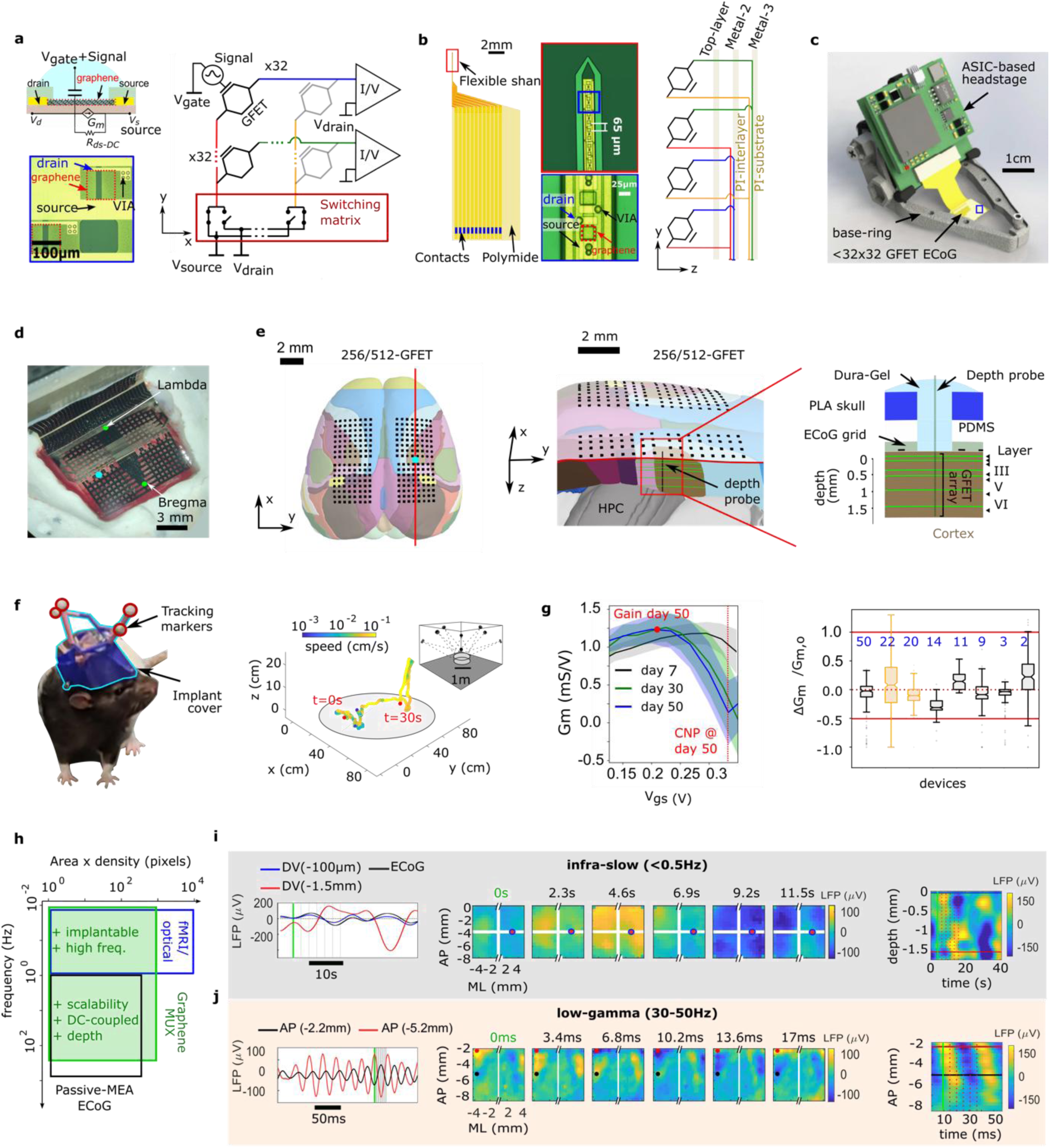
Graphene-based multiplexed large-scale recording technology. **a.** Schematic of a g-SGFET and its equivalent circuit (top-left), an optical micrography of the graphene transistor (bottom-left). In multiplexed g-SGFET array (right) each hexagon represents a g-SGFET, rows and columns are connected to the I-V converters and a switching matrix respectively. **b.** Layout (left), optical micrograph of the tip (middle) and schematic (right) of multiplexing of a 32-channel g-SGFET depth probe. **c.** The headstage and graphene ECoG assembled on a 3D-printed base ring for implantation. **d.** An example photo of the 256-channel g-SGFET ECoG array implanted over the neocortex of a rat. **e.** A schematics of top view of the ECoG array superimposed on the rendered anatomical map of the cortex (left) and zoom-in of the translaminar sagittal cut (middle) of the depth probe location. Surgical implant schematics (right) depicts ECoG grid and depth probe sealed by PDMS and dura-gel layers, sealed by skull replacement. **f.** Left: photograph of an implanted rat with enclosed miniature headstage and 3D-tracked markers during a freely moving recording. Right: an example 3D position and velocity (colorscale) trajectory of a rat in an open arena (multiple camera views used to triangulate rat 3D position and orientation, inset). **g.** Long-term stability of GFETs; Left: Voltage-dependent transconductance (Gm, median/std.dev.) for all GFETs in a 256-channel ECoG across days. Right: the change in Gm from the first day to the last day normalised by the Gm on day 1 for all functional GFETs in 256-channel (black) and 512-channel (orange) implanted ECoGs. Red lines, two-fold increase/decrease of Gm. Blue numbers, the number of days implanted for each device (see Supplemental Data Fig. 3). **h.** Schematic comparison of the spatio-temporal resolution of various imaging techniques. The advantages of multiplexed graphene probes compared to other techniques are indicated in green. **i.** Examples of traces in the infra-slow band (left) from representative ECoG and two intracortical channels. Infra-slow voltage maps recorded with a 256-channel ECoG (middle). The white gaps indicate gaps between channel rows/columns (see panel d,e). Red circle, location of the depth probe. IsLFP depth profile (right); blue and red lines, the channels shown on the left panel. **j.** Examples of LFP traces in the low-gamma band (left), and corresponding voltage maps (middle) and AL-PM section of profile (right) recorded with a 512-channel ECoG. The red and black dots/lines, LFP channels plotted on left.

The scalability of multiplexed graphene arrays has been theoretically supported^44,45^, but reproducible large-scale probe fabrication with high yield and detailed characterization of long-term performance *in vivo* has remained elusive. Here, we fabricated graphene transistor grids of 1024 (32×32), 512 (32×16) and 256 (16×16 or 32×8) channels and evaluated performance in terms of wide band sensitivity. Addition of columns reduces the load resistance at the input of the transimpedance amplifier, affecting the floor noise of the system (Supplemental Data Fig. 1). Below the 1/f corner frequency, transistors showed optimal performance with a median equivalent noise at the gate (*V*_*gs*−*rms*_) of 3.4µV (Q1= 2.8µV and Q3=6.0µV) per frequency decade for transistors of 100µm × 100µm (Supplemental Data Fig. 1). When considering a wide frequency-band (0.1-100Hz), we found that 32×16 (i.e. 512-channel probes) presented a good trade-off between channel count and sensitivity with a median *V*_*gs*−*rms*_ of 9.7µV (Q1= 8.1µV and Q3= 13.7µV) (Supplemental Data Fig. 1).

To overcome the limitation of the existing technology to 2-dimensional ECoG arrays and enabling volumetric sampling of the DC potential, we extended the concept of multiplexed arrays to depth neural probes by embedding addressable transistor arrays into high aspect-ratio flexible shanks. By expanding the second dimension of the array across metal layers in a metal-insulator stack (Fig. 1b), we fabricated a 32-channel, 106µm-wide, multiplexed, flexible probe based on 3 metal layers^30,47,48^. Depth probes showed a comparable median *V*_*gs*−*rms*_ of 5.3µV (Q1= 3.9 µV and Q3= 8.8 µV) consistent with the smaller transistor dimensions of 20µm x 30µm (width x length) (Supplemental Data Fig. 1). Importantly, owing to their shared multiplexing architecture, brain conformable ECoGs and depth probes could be seamlessly combined and operated with the same recording system (by sharing read-out blocks and sampling resources) with minimal connector overheads, enabling DC-coupled, dense recordings across cortical layers simultaneously with surface recordings. To enable naturalistic recordings, we further developed custom front-end electronics. We scaled custom open-source discrete electronics system^45^, and deployed a compact head-stage based on an application-specific integrated circuit (ASIC)^43^ to enable control and acquisition of, respectively, 16×16 and 32×16 channels of DC-coupled LFP in freely moving rats (Fig. 1c, Methods and Supplemental Data Fig. 2).

### Chronic implantation of combined large-scale ECoG and depth graphene probes

To explore the spatio-temporal dynamics of wide-band LFP during spontaneous behaviour, we developed a surgical approach enabling chronic implantation of the large-scale ECoGs to bilaterally cover a large fraction of the dorsal neocortex (ca. 12mm x 9 mm) in 8 rats (6 of them with 256-channel and 2 with 512-channel ECoGs, Fig. 1d-e, Methods). The surgery was split in two phases: implantation of a base-ring support structure exposing the full dorsal cranial surface (Fig. 1c), and implantation of the large-scale ECoG and depth probes following full recovery of the animals (Fig. 1d). After placing the ECoG on the brain surface, it was covered with a PDMS layer and a tightly sealed 3D-printed skull replacement with predesigned opening for the subsequent targeted insertion of the depth probe into the region of choice (somatosensory cortex, ML 3.2mm/ AP −-3.4mm, n=2). The insertion of the flexible depth probe through an opening in the ECoG was facilitated by a rigid silicon shuttle^48,49^ (Methods, Fig. 1e, right). Electrophysiological recordings were performed longitudinally over several days (10 to 50 days) in combination with high-precision, high-speed 3D motion capture of the rat’s head (Fig. 1f) for the classification of behavioural states in 32 recording sessions lasting between 46 min and 248 min (mean of 145.8 min). The chronically implanted neural probes showed stable sensitivity, quantified by transconductance (Gm) of transistors, for up to 50 days (see Fig. 1g and Supplemental Data Fig. 3).

The large-area coverage and high spatio-temporal resolution measurement of the DC-coupled LFP enabled by joint recording with ECoG and depth probes provided a unique 2.5D imaging-like electrophysiological window into DC-coupled brain dynamics (Fig. 1h), as illustrated by the examples of spatio-temporal LFP patterns in the wide frequency range from the infra-slow (Fig. 1i) to gamma band (Fig. 1j) and Movies 1 and 2, respectively. Surprisingly, isLFP exhibited rich spatio-temporal dynamics both across cortical areas and layers. In the following sections we demonstrate the technical advantages and power of the new technology in revealing that DC-potential shifts and isLFP dynamics arise from the spontaneous alternation of physiological brain states.

### Spatio-spectral framework for identification of brain states

Spectral content of the LFP or EEG is informative of the associated brain states, but requires complimentary motor-state measurements to disambiguate it well^50^. Wide-band topographic large scale LFP sampling enables extending single- or few-channel spectral features used for brain state classification^51,52^, to spatio-spectral features that would include anatomical information of the network dynamics defining each state leading to improved accuracy of detection. The same rationale underlies major advances made possible by high-density multichannel probes, which enable spike sorting of single units and the classification of high-frequency oscillations based on their invariant spatial sources^53,54^.

To compress joint spatio-spectral structure of the LFP, multitaper spectrograms were first integrated into spectral power within various frequency bands, from delta to low-gamma and the resulting spatio-spectral matrix reduced to the first three principal components. These features were further used for unsupervised brain state classification based on a Gaussian mixture model (GMM), enabling reliable identification of three major electrophysiological states: slow-wave (SW) state associated with large power slow wave activity, theta state associated with theta rhythm and high-voltage spindles (HVS)^55–57^ (Fig. 2a). The electrophysiological state segmentation was complemented with the motor state extracted from the 3D tracking data to subdivide the physiological states into slow-wave sleep (SWS), rapid eye movement (REM) sleep, awake non-theta (Aw. NTHE), awake theta (THE) and micro-arousals (MA - defined as short bouts of micromovement during SWS) (Fig. 2b and Methods). The extracted states presented unique spatio-spectral features across cortical regions (Supplemental Data Fig. 4), as exemplified by a spatially localized sigma power in anterior regions during SWS (Fig. 2c) and the presence of prominent peaks (at 8Hz and higher-order harmonics) aligned with the somatosensory cortex during HVS (Fig. 2d).

**Figure 2.**
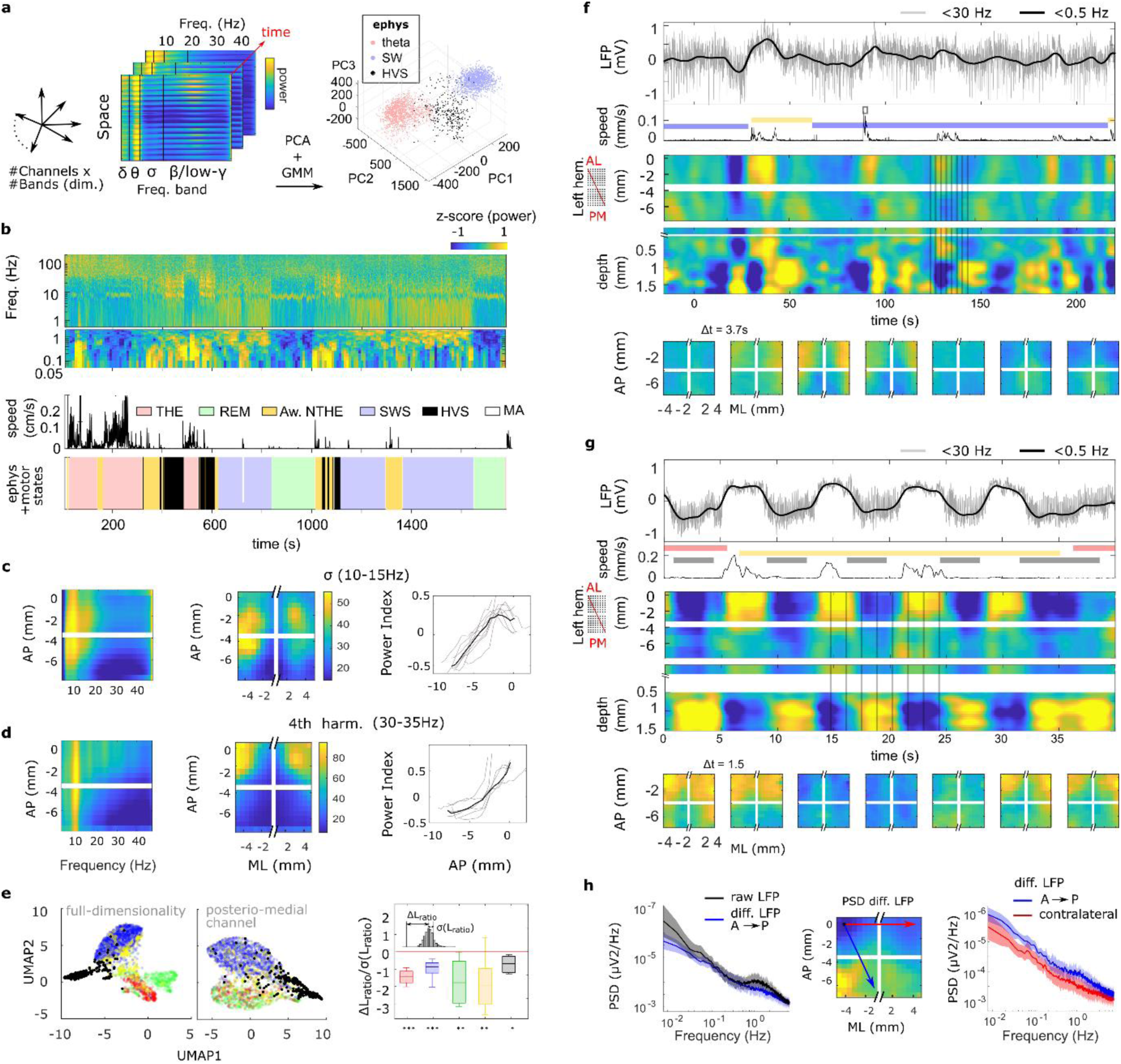
Spatio-spectral identification of the brain states. **a.** Description of the brain state classification procedure. Spectral content in various frequency bands in all channels is used for classification based on PCA compression and a GMM. **b.** From top to bottom: Example spectrogram spanning from infra-slow (0.05Hz) to high-gamma (200Hz) bands, speed of the animal and brain state obtained from the combination of electrophysiological and motor states. **c.** Spatio-spectral characteristics of SWS. Mean spectrum (ML-axis-averaged) across the AP axis (left). Mean spectral power map within a narrow band (top labels) across the array (middle); group summary of the power index (see Methods) across the AP axis (right). Gray, single animals; black, group mean (n=8/6 for SWS/HVS). **d.** Same as c for HVS. **e.** UMAP embedding of spatio-spectral characteristics from all channels (left) and from an example posterior channel (middle); right, separability of individual brain states measured as normalized difference of *L_ratio* between using all channels and the median of that using individual channels (n=8 for THE, SWS and Aw. NTHE; n=6 for REM, n=5 for HVS, one-tailed Wilcoxon signed-rank test p-value: *<0.05, **<0.02, ***<0.01). **f.** From top to bottom: LFP in an anterolateral channel of the ECoG during a long SWS epoch interrupted by micro-arousals and sub-threshold micro-movements (<30Hz, gray and <0.5Hz, black). Speed of the animal overlaid with the states colored as in b. Color-coded isLFP profile from the anterolateral corner to the posteromedial end of the ECoG for the left hemisphere. Depth profile of the LFP (above zero, the isLFP in the ECoG channel nearest to the depth probe is shown). The vertical lines indicate the timestamps of the ECoG isLFP frames shown below. **g.** Same as in e, but during an epoch containing multiple consecutive HVS events. **h.** Mean PSD during immobility (wake and sleep) for the raw LFP and the LFP re-referenced to an anterolateral channel (left, n=6). Map of the re-referenced LFP power in the 0.01-0.1Hz band (middle). The arrows indicate the two pairs of channels used to compute the power of the re-referenced LFP in the right panel. Comparison of the power of the re-referenced frontal LFP to posterior and contralateral positions as shown by arrows in the middle panel (right, n=6).

Using the obtained state labels as ground truth, we assessed how spatially structured spectral content of the surface LFP contributed to the state-separability without considering the motor information. In contrast to single channel spectra, the full spatio-spectral data provided significantly better quality of state separation quantified using cluster isolation metric (*L_ratio*) in the Uniform Manifold Approximation and Projection (UMAP) embedding of spatio-spectral vectors (Fig. 2e, see Methods). Spatio-spectral structure enabled segregation of states that are otherwise spectrally strongly overlapping, such as THE and REM (Fig. 2e; one-tailed Wilcoxon signed-rank test).

### Brain state alternation co-occurs with topographic and translaminar DC-potential shifts

Could state transitions explain isLFP dynamics? Indeed, isLFP exhibited prominent patterns with spatial and translaminar phase reversals tightly correlated with brain state transitions (Fig. 2f,g). During SWS, surface isLFP often presented a prominent anterolateral gradient linked to periods of increased arousal (Fig. 2f) and the phase reversal of isLFP around the cortical layer IV. Interestingly, a comparable pattern was observed during hypersynchronous HVS events, which were associated with a negative isLFP fluctuation, more pronounced in the anterolateral primary somatosensory cortex, and showed a similar translaminar phase reversal (Fig. 2g). To validate the consistency of spatio-temporal structure of the isLFP across animals we computed the LFP spectra during periods of immobility (Fig. 2h, left and Methods), reflecting both infra-slow power and a prominent band associated with UP-DOWN states alternation at ∼1-5 Hz^58^. In contrast, LFP spatial gradient, measured as LFP re-referenced to an anterolateral channel (dLFP), demonstrated a power-low shape with infraslow power spatially structured only in sagittal direction resulting from high coherence between symmetrical contralateral areas (Fig. 2h).

In order to identify and interpret the sources of these isLFP patterns we developed an analytical and physiological framework described in the following sections for distinct brain states.

### DC-coupled blind source separation: spike-and-wave discharges present topographically and translaminarly matching DC shifts

HVS state is defined by hypersynchronous thalamocortical oscillations, or spike-and-wave discharges, and is considered a model of absence epilepsy^56,59^. HVSs have been linked to spatially inhomogeneous DC shifts^42^ and hence represents an excellent frequently occurring (Fig. 3a) benchmark physiological pattern for dissecting the DC-potential origin of the isLFP and guide the discovery of biomarkers for the detection and localization of the epileptic foci^27,60,61^.

**Figure 3.**
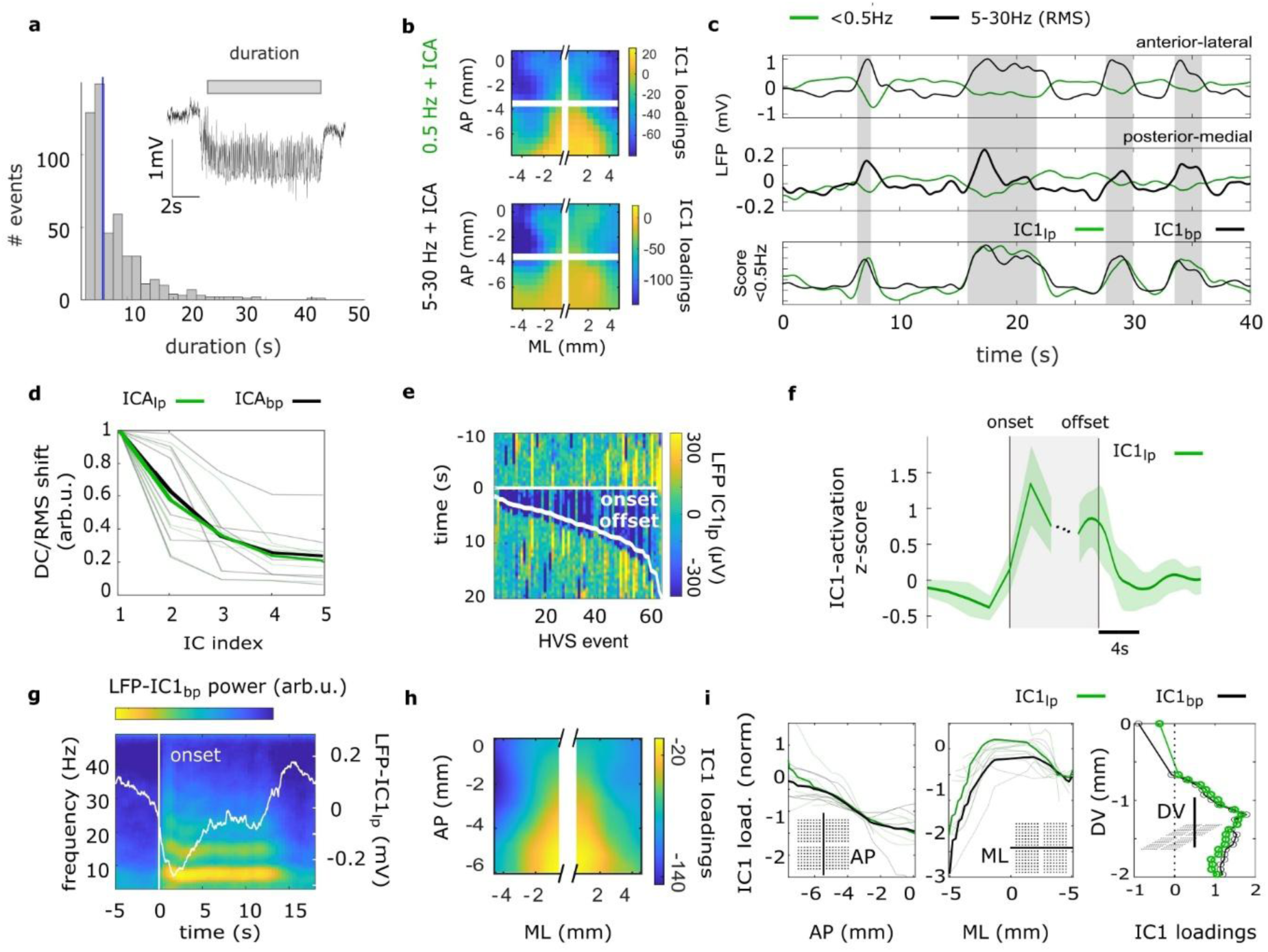
Topographic structure of infra-slow and spike-and-wave sources of high voltage spindles. **a.** Distribution of HVS duration across animals (median, blue line). Inset, an example of full-band LFP during HVS. **b,** IC1 loadings extracted from the isLFP (IC_lp_, top) or spike-and-wave band (5-30Hz) (IC_pb_, bottom). **c**. Representative traces of LFP_lp_ and RMS of the LFP_bp_ at an anterolateral (top) and posterior-medial (middle) channels and activations for IC1_lp_ and RMS of the IC1_bp_ (bottom). **d.** Mean DC-potential shift for IC_lp_ and RMS change for IC1_pb_ around the HVS onset (±2s) for the first five ICs (see Methods, n=6). **e.** Colour-coded reconstructed LFP from the IC1_lp_ across HVS events for an example animal. White lines indicate state onset and offset. **f.** Group (n=6) average IC1_lp_ score triggered on the onset and offset of the detected HVS events. Shading, std. dev. **g.** The triggered average spectrogram of the reconstructed LFP from the IC1_bp_ around the HVS onset overlaid with the triggered average of the infra-slow reconstructed LFP for the IC1_lp_ for an example animal. **h.** Map of group median loading of the IC1 (n=4 animals with consistent coverage). **i.** Comparison of spatial profiles of IC1_lp_ (green) and IC1_bp_ (black) loadings across animals (n=6) in the three axes (thick line shows the mean across animals). IC1 loadings were normalized by their value at −3.5mm w.r.t. Bregma in the AP axis, 4mm w.r.t. the midline in the ML axis and at −1mm w.r.t. the surface for the DV axis.

To reveal and study the relation between topographic and translaminar sources for spike-and-wave activity and DC-potential shifts, we developed an analytical approach based on joint blind source separation of both spatio-temporal patterns in the of infra-slow band (low-pass below 0.5Hz: lpLFP) and broad-band spike-and-wave bands (band-pass of 5-30Hz: bpLFP). Independent component analysis (ICA) applied to lpLFP and bpLFP centred on HVS epochs identifies independent temporal dynamics, and associated spatial maps, of the main sources for DC-potential shifts (IC_lp_) and spike-and-wave discharges (IC_bp_). Interestingly, the first IC (IC1) extracted from the two bands showed comparable topographic structure (Fig. 3b) and temporal dynamics (Fig. 3c) between the two bands, revealing tight spatiotemporal correlation of the LFP sources in the two bands. Furthermore, the magnitude of the IC_lp_ and IC_bp_ showed comparable relative attenuation across sorted ICs (Fig. 3d), indicating DC-potential shifts and changes in oscillatory power are similarly explained by a reduced number of components. The activation of IC1_lp_ around the HVS onset (and offsets) confirmed that topographic DC-potential shifts and return to baseline were closely aligned with the synchronous state activation (Fig. 3e,f). This match in topographic structure of the IC_1_ loadings between the two bands was consistent in all animals (Fig. 3h,i) and was also present across cortical layers with a phase reversal around layer IV (Fig. 3i).

Overall, these results demonstrate temporally, topographically and translaminarly aligned sources for localised spike-and-wave discharges and DC-potential shifts, suggesting a direct relationship between the physiological mechanisms generating these wide-band dynamics.

### Spatiotemporal isLFP decomposition across sleep sub-states links DC-potential shifts to synchrony fluctuations

Recent studies have explored the relationship between the level of arousal and the microstructure of sleep sub-states^62–64^. During SWS, the power of sleep spindles varies at the infra-slow time scale, marking cycles in sleep fragility^62^ linked to neuromodulatory dynamics^7,65–67^. Inferring sleep microstructure from isLFP correlates of these neuromodulatory and thalamocortical synchronisation dynamics would provide a powerful biomarker, but it remains poorly studied due to limited spatial sampling, lack of stable DC-coupled recordings across unconstrained behavior and poor interpretability of the DC signals^26,31,32,68,69^.

To characterize the spatial structure of isLFP associated with sleep sub-state transitions we analysed long sleep sessions including many cycles of SWS, MA and REM sleep (Fig. 4a) and extracted spatial patterns using singular value decomposition (SVD) of the isLFP (see Methods). While the 1^st^ SV component (SV1) contained a trivial global variation in the signal amplitude, the 2^nd^ component (SV2) reliably displayed an anteroposterior-mediolateral sign-switching gradient that was topographically consistent across animals (Fig. 4b). The SV2 score, reflecting the temporal dynamics of this topographical isLFP pattern, presented prominent troughs that temporally correlated with the onsets of MA and REM epochs (Fig. 4c,d), suggesting a correlation between this isLFP component and the depth of sleep. The inter-event interval between SV2-score troughs ranged between 10 and 100 seconds, with the median 45.2 across animals (Fig. 4e). This distribution is consistent with previous reports of neuromodulatory dynamics^8^, and implies the detection of covert MA (Fig. 4e)^7^. In turn, the amplitude of the SV2-score troughs correlated with the integrated micro-motion of the animals around covert MA (Spearman correlation r=0.22, p-value = 0.0017, Supplemental Data Fig. 5).

**Figure 4.**
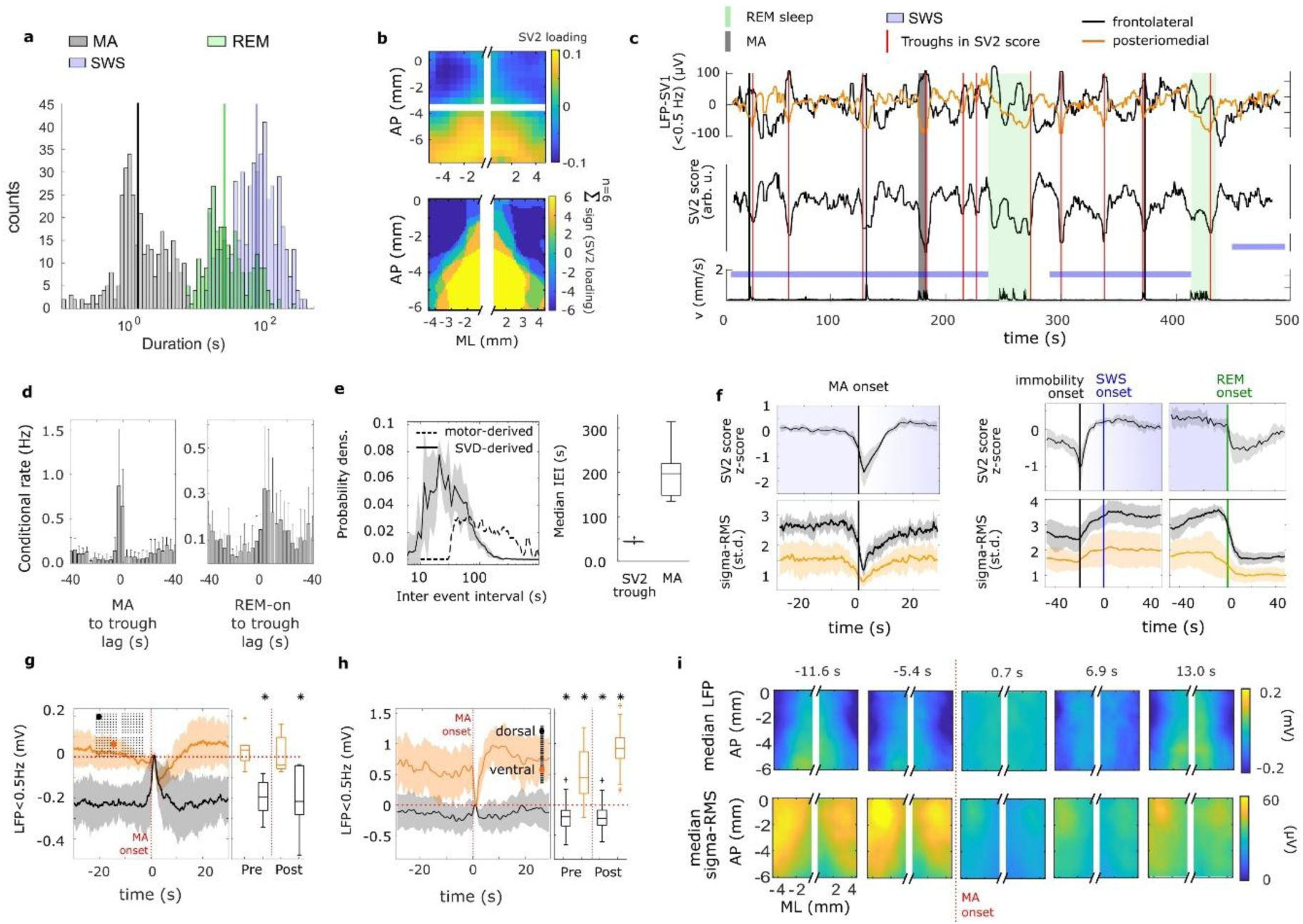
isLFP correlates of sleep state transitions. **a.** Duration of detected REM sleep, SWS and MA epochs across animals. **b.** Color-coded spatial map of the SV2 loadings for an example animal (top) and spatial map of the cross-animal (n=6) consistency of the SV2 loadings (see Methods, bottom). **c.** From top to bottom: LFP after subtracting the contribution from SV1 for two reference channels, score of the SV2 (<30Hz) and (<0.5Hz), brain states and velocity of the animal head. Red lines, troughs of SV2 score. **d.** Group mean cross-correlogram between state onsets and the SV2 score troughs (standard deviation as error bars, n=6 animals). **e.** Left: distribution of inter-event intervals (IEI) between SV2 score troughs during SWS, mean and std.dev. across animals (n=6) and the IEI distribution between MA accumulated across 6 animals. Right: boxplot of median IEI for SV2 score troughs and MA across animals (n=6 animals). **f.** Top, triggered average of the SV2-score around the MA onset (left) and at the onset of SWS and its transition into REM (right). Bottom, corresponding average normalized spindle power for anterior (black) and posterior (orange) reference channels (n=6, shading is 1 standard deviation). **g.** Left, Average isLFP aligned to the MA onset and referenced to the aroused state, for two positions (mapped on the array in the insets, red, black colors) on the surface across animals (left, n=6). Thick lines, mean; shading, std.dev. Right, boxplot of mean isLFP across animals at times −10s and 10s around the MA onset (Wilcoxon signed-rank test p< 0.05). **h.** same as g for one animal and in depth across events (39 events). Superficial channel (DV = −100μm) in black and deep channel (DV = −-1.1mm) in orange. Pre and post times in the boxplot correspond to −10s and 10s around the MA onset (Wilcoxon signed-rank test p< 0.05). **i.** Median topographic maps of the isLFP re-referenced to MA state (top) and sigma power (bottom) aligned around the SWS-MA transitions (n=6).

The SV2-score fluctuations around MAs suggest the sequential de-activation and re-activation of a SWS-related topographic DC-potential (Fig. 4f, top left). This activation was also evident at the onset of SWS and was followed by an abrupt de-activation at the transition into REM sleep (Fig. 4f, top right). These SV2-score dynamics mirrored the sigma-power dynamics, which showed a similar gradual onset into the SWS and abrupt offset from it (Fig. 4f, bottom and Supplemental Data Fig. 6). Analogously to the HVS case, the negative DC shift in anterolateral areas associated with SWS was related to a larger sigma power in anterolateral areas (Fig. 4f). These observations suggest that, similar to HVS, SWS state defined by synchronous slow oscillations and sleep spindles is associated with a DC shift with the similar anteroposterior gradient.

### Desynchronized-state referencing: interpreting DC polarity across sleep sub-states

The mapping of isLFP poses a challenge to interpret the polarity of topographical DC-shifts^70,33^, with the absolute DC-potentials arbitrarily defined by the electrochemical potential between the graphene channel and the reference electrode.

Following the apparent DC-potential sources activation linked to high synchrony state onset, we propose to refer the DC-potential to the desynchronised states to interpret the polarity and magnitude of the DC-potential shifts associated to SWS. Indeed, referencing the isLFP (<0.5Hz) to the median DC-potential during MA and awake states revealed a negative DC shift reaching, on average, the order of 200 µV in surface derivations over anterolateral areas during the SWS state (Fig. 4g). Similarly, the translaminar isLFP around MA referenced to the aroused state also showed the phase reversal of the DC potential around layer IV with abrupt onset of deep positive shifts in the 0.5-1 mV range upon transition upon the SWS onset (Fig. 4h). As predicted by SV2 loading (Fig. 4b), the topographic structure of the DC shifts in the surface LFP and spindle power showed consistent temporal progression through SWS to MA transitions (Fig. 4i and Supplemental Data Fig. 6).

Taken together, these results show that the isLFP exhibits topographical and translaminar dynamics reflecting transitions to and from the thalamocortical synchronization characteristic of SWS. By referencing the DC potential to the desynchronized state, the polarity of these shifts can be unambiguously interpreted as synchrony-related, closely resembling those associated with HVS.

### Decomposition of transient spatio-spectral states: detection of local sleep spindle generators

These results highlight the need to directly examine how individual sleep spindles are topographically coupled to local excitability fluctuations reflected in the isLFP and extending the analysis to the next slow oscillation time scale (∼1Hz) reflected by slow waves associated with UP/DOWN state transitions^71–73^. To address this question and demonstrate the power of large-scale electrophysiological imaging to identify local oscillatory states we set out to detect local generators of sleep spindles.

Multiplexed graphene probes provided smooth maps of sigma band (10-20Hz) power with striking topographically structured dynamic patterns that presumably reflect the activation of various cortico-thalamic networks involved in the generation of sleep spindles (see [Movie 3](https://gin.g-node.org/RGCortadella/LFP-imaging/src/master/Movie_3.avi)). To characterize the dynamics in sigma band with both fine spatial and spectral resolution, we decomposed the power spectrograms across channels and frequency bins using factor analysis^52^ (see Methods and Supplemental Data Fig. 7). The spatio-spectral structure of each of the factor loadings reflected diverse spatial sources of spindle oscillations (Fig. 5a) with their respective factor score dynamic displaying transient bouts corresponding to waxing and waning oscillations in the anatomical location and at the specific spindle frequency reflected by the respective factor loadings (Fig. 5b). The dynamics of these factors represent a lower-dimensional description of the spatio-temporal variation of sigma power as illustrated in Fig. 5c. Group statistics revealed comparable factors across animals (Fig. 5d and Supplemental Data Fig. 7) with stereotypical spindle burst durations (0.7±0.2s across animals and factors, Fig. 5e) and harmonic index, a measure of spectral concentration (0.7±0.8, across factors and animals, Supplemental Data Fig. 7 and Methods). Spindles in anterior areas (Fig. 5d) present larger amplitude compared to spindles in posterior areas (Fig. 5d,f).

**Figure 5.**
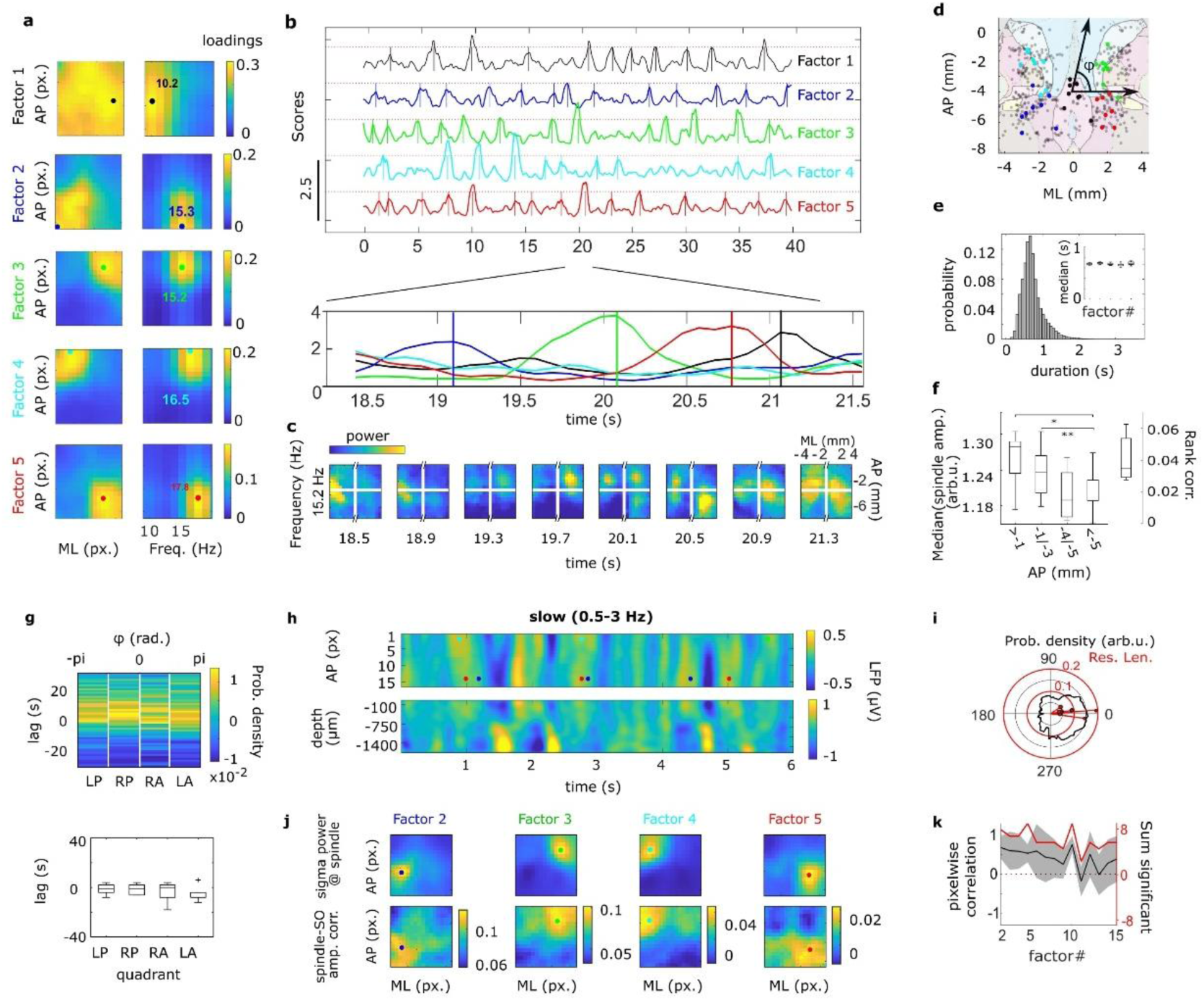
Detection of diverse local sleep spindle generators from the cortical surface. **a.** Spatio-spectral loadings of the first five factors for an example animal. Filled circles mark the position in space and frequency of the maximum loading. **b.** Score dynamics for the first five factors in panel a. The bottom plot shows overlapping traces in a zoomed-in window. **c.** Example of power dynamics at 15.2Hz across frames corresponding to the time window shown in b. **d.** Position of loadings peak in space (first 30 factors per animal). Coloured dots correspond to matching factors (n= 8 animals). **e.** Distribution of half-maximum width of the score peaks across the first five example factors. The inset describes the median of sleep spindle duration (in seconds) for all matched factors across animals (n=8 animals). **f.** Median spindle power for all factors localized across the AP axis (n = 8 animals). Wilcoxon signed-rank test, * p-val=0.031, ** p-val=0.016. The right-axis shows the rank correlation between amplitude of spindles and the AP-position of their maximum loading across animals. **g.** Cross-correlogram between the troughs in the AL isLFP (<0.05 Hz) and time of occurrence of spindles detected in different quadrants (top). The time lag of maximal cross-correlation across animals and ECoG quadrants (bottom). No significant differences observed between different quadrants (paired-sample t-test, n=6). **h.** An example of the LFP in the 0.5 - 3 Hz band for the 16 channels in the fourth column of the ECoG array from the left hemisphere (top) during SWS. The coloured dots correspond to the time stamps for the spindles detected with factors 2 to 5. **i.** Phase distribution of SO at the time of all spindles (for factors 1–10) across animals (n=8). In red, the resultant vector for the phase of SO at the time of spindles for 8 animals. **j.** Top: maps of mean spindle band power at the detected spindle time points for an example animal for factors 2 to 5 (same as in Fig. 5a and 5b). Bottom: maps of the correlation between the amplitude of the SO cycle and the UP-locked spindle power for factors 2 to 5 (same as in Fig. 5c and 5d). **k.** Correlation of the spindle power maps and the SO-spindle correlation maps in the hemisphere of maximum loading across 15 factors sorted by variance (mean, shaded standard deviation). The right axis (red) shows the sum of significantly correlated factors across animals for the first 15 factors (permutation test p<0.05, 1000 permutations).

### Large-scale cross-frequency coupling: global infra-slow and local slow modulation of sleep spindles

Next, we investigated the spatial and temporal relationship between local sleep spindles and the spatial structure of isLFP fluctuations within long stretches of SWS uninterrupted by overt MA (Supplemental Data Fig. 8 and [Movie 4](https://gin.g-node.org/RGCortadella/LFP-imaging/src/master/Movie_4.avi)). Interestingly, independent of their localization, sleep spindles are temporally correlated to the troughs of isLFP at the anterior-lateral regions (Fig.5g). The fact that infra-slow fluctuations globally modulate local sleep spindles suggests that, although the detected spindle generators are orthogonal and present a low degree of overlap at a faster time scale (Fig. 5b), their rate is strongly co-modulated by the infra-slow dynamics reflected by isLFP fluctuations (Supplemental Data Fig. 9). These results suggest that the gradient in isLFP likely reflects the distinct power of spindles along the AP axis.

At a faster time-scale, SO dynamics are known to entrain sleep spindles^71,72^. However, while local slow waves and sleep spindles have been reported before^74,75^, their spatio-temporal relationship is not known. Leveraging large-scale, dense ECoG recordings we detected both local and global SO (with the surface negative delta waves reflecting the characteristic phase reversal around L-IV^76^, Supplemental Data Fig. 10) and related their dynamics to that of sleep spindles (Fig. 5h). Spindles bursts, regardless of their origin, were locked to the peak of global surface slow waves, consistent with their occurrence during the Up states^77^ (Fig. 5i). Strikingly, the trough-to-peak amplitude of the local SO (Fig. 5i) correlated with the power of co-occurring sleep spindles in a topographic manner (Fig. 5j,k). The fact that correlation maps reached their maximum at the location of the co-occurring spindles indicates the tight spatio-temporal relationship between local DOWN-UP states and sleep spindles, with both their timing and the amplitude of the local sleep spindle oscillations correlating to that of the local UP-state (see Supplemental Data Fig. 11). Altogether, these results illustrate the power of large-scale, high density, broad band LFP mapping for dissecting and interrelating rich SWS dynamics across cortical regions and temporal scales.

### [K⁺]ₒ gradient–driven dipoles: a mechanistic framework for interpreting the isLFP

Despite the long history of DC and infra-slow potential measurements^23,26,32^ the biophysical source of infra-slow field potentials remains unclear, with some works suggesting a non-neuronal origin^5,26^. Our characterization of synchronous oscillatory network dynamics and isLFP across cortical regions and layers yields new constraints on the possible origin of isLFP, providing a framework for the biophysical interpretation of the DC-coupled recordings.

Our results show that transitions from synchronous to desynchronized states were associated with spatially structured DC-potential shifts. Both the spike-and-wave discharge component of HVS and the associated DC-potential shift had topographically identical sources centred on somatosensory areas (Fig. 6a, left and Fig. 3h,i). Similarly, the topography of DC shifts associated with SWS state, presented a gradient from antero-lateral to posterior-medial areas (Fig. 6a, right, Fig. 4, Supplemental Data Fig. 8) matching the topography of sleep spindle power (Fig. 6a and Fig. 2c, 4i, 5f). Furthermore, HVS-associated translaminar DC profile had a phase reversal around layer IV, similar to that of spike-and-wave discharge^78^ (Fig. 6b, left and Fig. 3), and infra-slow fluctuations linked to SWS showed a translaminar profile comparable to that of delta waves^79,80^ (Fig. 6b, right).

**Figure 6.**
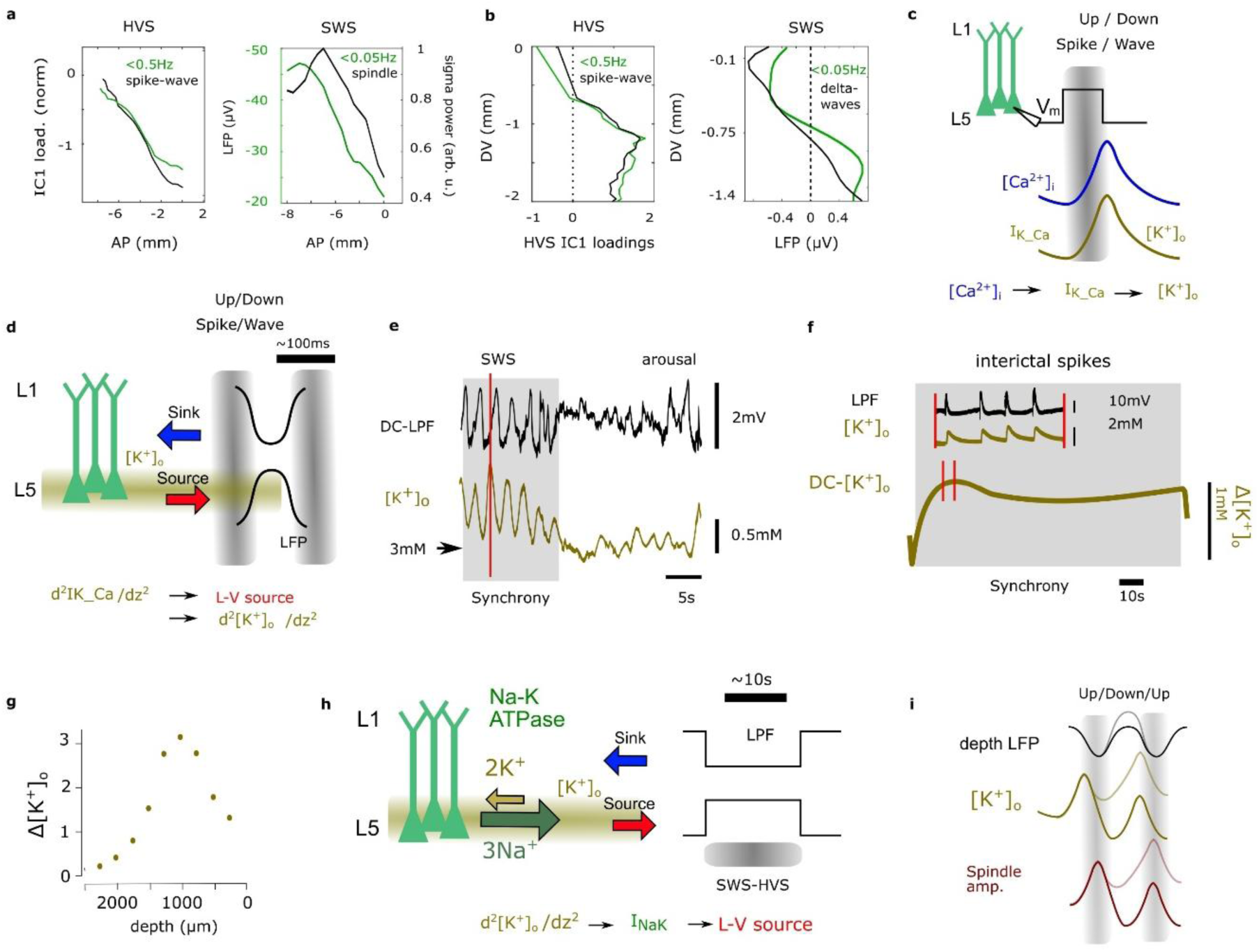
Qualitative model of the link between synchronous states and DC-potential changes. **a.** Topographical relation between the isLFP amplitude profile and amplitude of spike-and-wave discharges (left) and the power of sleep spindles during SWS (right). **b.** Translaminar profile of isLFP and the HVS spike-and-wave amplitude (left) and slow wave profile (right). **c.** Synchronous states such as UP-states and the spike component of HVS drive an increase in intracellular calcium, which leads to the opening of *K* channels that hyperpolarize the neurons. **d.** The increase in *K* conductance during the UP-DOWN-state transition is larger in layers with high neuronal activity, leading to an LFP source around L-V and generation of a *K*^+^ gradient around this layer. **e.** Illustration for the relation between extracellular potassium concentration and slow oscillations as well as its DC level change between SWS and desynchronized states (adapted with permission from Seigneur et. al. 2006^97^). **f, g.** Experimental data showing the buildup of [ *K*^+^]_*o*_during an induced seizure (f) and its depth profile (g) (adapted with permission from Moody et. al. 1976^99^). **h.** The buildup of a layer-specific DC-level of [*K*^+^]_*o*_ leads to a layer-specific activation of the electrogenic Na-K pump, which in turn generates a DC current source giving rise to isLFP. **i.** Schematic of the relation between the amplitude of slow oscillations, potentially generated by changes in the potassium conductance during the DOWN-state and the accumulation of potassium in subsequent cycles leading to increased excitability and sleep spindle amplitude.

Common to all synchronous cortical dynamics, the active phase (spike phase of the HVS or the UP-state of the slow oscillation) is associated with synchronous neural firing resulting in an increase of intracellular *Ca*^2+^ concentration^81,82^. This leads to the activation of multiple *K^+^* currents, most prominently, calcium-dependent potassium (*I*_*K*−*Ca*_), as well as ATP-dependent *K^+^* and inward-rectifier *K^+^* currents, that are responsible for the termination of the active state, repolarization of the membrane potential and an increase in extracellular potassium concentration ([*K*^+^]_*o*_)^83–88,82,89–92^ (Fig. 6c). These activity-dependent hyperpolarizing currents are larger in layer V, where there is higher recurrent excitation and firing rate^93–95^. This implies a higher net *I*_*K*−*Ca*_ localized in this layer during the DOWN-state-associated delta wave^80,96^ and HVS waves^78^ (Fig. 6d), leading to a layer-specific accumulation of [*K*^+^]_*o*_. During the wave phase of HVS or the DOWN-state, the extracellular potassium ions are buffered by glia^97^ and pumped back into the cell by NaK-ATPase pump^98^. However, under repeated activation of *K^+^*conductance during synchronous activity, these mechanisms are not sufficient to return the [*K*^+^]_*o*_ to baseline level. This results in a local build-up of [*K*^+^]_*o*_ ^99–103^, leading to a state-dependent concentration gradient *Δ*[*K*^+^]_*o*_ (Fig. 6e,f)^97,99,102^, with a sustained layer V-dominance^99,102^ (Fig. 6g). This layer specificity is expected to match the delta/HVS wave profile (Fig 7b) that originated the gradient in [*K*^+^]_*o*_.

Here, we propose that this sustained [*K*^+^]_*o*_ gradient during states of high synchrony will lead to the generation of a sustained electrical dipole by activating the electrogenic NaK-ATPase pump and, correspondingly, an outward current with amplitude dependent on a translaminar concentration^98,103^ (Fig. 6g, h). In this manner, the sustained and layer-V localized concentration gradient of [*K*^+^]_*o*_ would be a consequence of neuronal synchrony giving rise to a similar depth-reversing DC dipole as that of the delta/HVS wave that gave rise to the *K^+^* accumulation (Fig. 6h). Importantly, the mechanism we propose here is independent of the source of synchrony, and is consistent with numerous prior work showing that both electrical stimulation of the neocortex or spontaneous and induced seizures results in both [*K*^+^]_*o*_ accumulation with matching translaminar gradient as well as translaminar reversal of the extracellular DC potential from negative at the surface to positive in deep layers^23,26,30,99,101,102,104^.

Activation of the outward current driven by the NaK-ATPase pump at the same time, could causally influence the network dynamics by increasing the resting potential of neurons and contribute to terminating a synchronous state^103^. This causality could be reflected on the finer timescale in the relation between the larger amplitude of delta waves associated with a larger increase in [*K*^+^]_*o*_ and higher excitability during the following UP-state as reflected in the amplitude of sleep spindles (Fig. 6i).

## Discussion

### Multiplexed graphene transistor arrays enable DC-coupled, 2.5D electrophysiological imaging

Here, we establish a scalable electrophysiological platform based on multiplexed graphene transistor arrays that enables large-scale, DC-coupled, epicortical and intracortical LFP recordings in freely moving rats. While previous advances in flexible micro-ECoG arrays enabled increased sampling densities across large cortical areas^37–39^, non-multiplexed designs were constrained by connector complexity and front-end electronics, limiting scalability and miniaturization^38,39^. Conversely, multiplexed flexible arrays with high channel counts have so far lacked the technological maturity for their application in chronic settings in freely moving animals^36^. The presented technology overcomes these limitations and enables brain-wide, 2.5-dimensional DC-coupled electrophysiological imaging, providing simultaneous access to infra-slow and fast network dynamics across space, depth, and behavioral state.

Leveraging these technical developments, we uncovered the link between synchronous brain states and the spatio-temporal and translaminar structure of the isLFP. Although the biophysical origin of isLFP remains debated and likely reflects multiple contributing processes^23,26,61,5^, the consistent coupling we observed between synchronous neural states and structured DC patterns substantially constrains possible mechanisms. The infra-slow dynamics we identified in the isLFP are consistent with that observed in fMRI^11,17,105^, neuromodulatory signals^7,66^ and slow fluctuations of oscillatory power^12,62,106^, supporting the interpretation that isLFPs reflect spatially organized changes in cortical excitability.

### Large-scale spatio-spectral brain state organization

We demonstrated the utility of dense large-scale recordings by identifying synchronous brain states in an unsupervised manner based on the spatial structure of LFP spectral maps. Unlike conventional single-channel approaches, large-scale topographic LFP sampling leverages the state-specific spatial synchronization across cortical and subcortical regions, improving sensitivity to cross-scale dynamics and source localization^50,52,107^. Similar to spike sorting, large-scale dense recordings enable separation of overlapping spatiotemporal motifs that would otherwise remain conflated^4,108^. This approach is particularly relevant for capturing non-stationarity of brain dynamics associated with sleep sub-states such as local sleep^73,75^ or tonic-phasic REM alternations ^12,109,110^. The spatial structure of the HVS reported here is consistent with prior work in older rats^42,111^, while our higher-density recordings enabled detection of less synchronous, developmentally relevant oscillatory states characteristic of younger animals^59^. The detailed characterization of spatio-spectral features of brain states and identification of isLFP sources opens new avenues for diagnostics and therapeutical interventions via closed-loop electrical stimulation therapies based on biomarkers derived from wide-band LFP^26,61,112–116^.

### Multiscale intreaction linking spindles, slow oscillations, and infra-slow states

At a finer scale, we leveraged dense recordings to identify spatially localized and spectrally distinct sleep spindle generators. Regional diversity in sleep spindle frequency has been reported previously^117–122^, likely reflecting synchronization in distinct thalamocortical circuits. Our results provide an unbiased, data-driven confirmation of this organization, showing that slower spindles (∼10 Hz) tend to be more global, whereas most spindle events span a broad sigma–beta range and remain spatially localized. Spatio-spectral localization provides a sensitive and unbiased analysis of transient oscillations, analogous to the analysis of high-frequency oscillations, enabling the dissection of distinct circuit motifs and their functional investigation^52,108,123^.

Our results reveal a hierarchical relationship between infra-slow states and faster sleep rhythms, characterized by both emergent and reciprocal interactions. At the global scale, infra-slow states captured by the isLFP reflect slow fluctuations in the probability, synchrony, and spatial extent of slow oscillations and sleep spindles. The antero-posterior gradient of synchronization reflected in the isLFP topography was associated with the higher power of sleep spindles as well as slow waves, consistent with a higher probability of K-complexes originating in frontal regions^74^. At a finer spatial and temporal scale, however, SOs and spindles exhibit a more local and reciprocal relationship. Local UP states both emerge from and promote the occurrence of spatially aligned sleep spindles, as reflected by the strong correlation between local SO amplitude and spindle power. This coupling suggests that while infra-slow states provide a global excitability backdrop, local DOWN–UP transitions actively recruit spindle-generating circuits, giving rise to transient, spatially confined events such as K-complexes^72^. The precise directionality of causation, whether local UP states primarily trigger spindles or whether spindle activity feeds back to reinforce UP states, remains unresolved.

The precise detection and localization of sleep spindles and UP/DOWN-states could provide further insights into the mechanism by which these dynamics facilitate memory consolidation in distinct cortical areas^73,124,125^ with local packets of cortical synchrony differentially coupled with SPW-ripples^126,127^. Moreover, the heterogeneous content of the hippocampal replay within the infra-slow rhythm could be shaped by the spatio-temporal structure of the isLFP^46^, providing insights into the role of the region-specific sleep microstructure on memory cosolidation.

### Relationship between isLFP, neuromodulatory, [K^+^]_o_ and vasomotor dynamics

Neuromodulatory dynamics during sleep fluctuates at the infra-slow time scale of 0.01-0.05Hz (often referred to as ultra-slow^50^) and covaries with sleep spindle rate^84,63,62,65,7,66,128^. Neuromodulatory states that shape synchronous dynamics could, as we have shown, be directly inferred from the spatio-temporal isLFP. In fact, we show that isLFP dynamics can uncover states of increased arousal level reflected in extremely subtle (<0.5mm/s) movements during SWS, consistent with the relation of NE dynamics to overt and covert MA^7,128^.

While the origin of isLFPs is likely multifactorial, our results constrain viable mechanisms and support a testable model in which synchronous neuronal activity generates sustained, layer-specific extracellular K⁺ gradients, producing DC potential shifts. This framework resolves the longstanding ambiguity of DC polarity during synchronous states^23,31,26,61^ and motivates referencing DC signals to desynchronized states (REM, active wake) for consistent interpretation. Recently, neuromodulatory dynamics and brain state alternation have been shown to be bi-directionally causally interrelated to both systemic and local [*K*^+^]_*o*_^129,130^. High neuromodulatory levels are associated with elevated [*K*^+^]_*o*_, while it tonically decreases during NREM sleep and immobility. This low baseline sets up the backdrop for a strong driving force of outward *K^+^*, necessary for effective negative feedback in slow oscillations, and allows for large translaminar gradients of [*K*^+^]_*o*_ leading to a sustained dipole. Interestingly, increased [*K*^+^]_*o*_ also modulates vasomotor activity by the inward-rectifying channels in the blood vessels^5^. Therefore, our model supports that the laminar-specific changes in [*K*^+^]_*o*_ that generate sustained dipoles could also contribute to the delayed vasomotor dynamics. This could be the mechanism linking topographic infra-slow potentials measured by full-band EEG and resting state networks and the BOLD signals^21,105^.

### Conclusion

Together, the developed technology and analytical framework position DC-coupled, large-scale electrophysiology as a powerful complementary modality to functional imaging, providing a direct readout of infra-slow activity with high temporal resolution in freely behaving conditions. This approach opens new avenues for studying state-dependent coordination across cortical networks and for linking infra-slow dynamics to faster oscillations, with promising applications in both clinical and fundamental neuroscience.

## Methods

### Graphene probes microfabrication

Large-scale graphene neural probes were fabricated on a 4” wafer following the procedure described before^44^ with minor modifications. An initial 4µm-thick polyimide (PI-2611, HD MicroSystems) was spun-coated on a Si/SiO_2_ 4” wafer and baked at 350°*C*. Polyimide was chosen as a substrate due to its stability and biocompatibility^40,48,131,132^. A first metal layer 30nm Ti for adhesion, 300nm Au for conductance and 10nm Ni as diffusion barrier at high temperatures) was deposited by electron-beam vapour on a previously photodefined negative AZ 5412E (Clariant, Germany) and then structured by a lift-off process. Then, a 2µm-thick polyimide interlayer was deposited to separate metal layers and vertical interconnection access (VIA) holes opened using a 200nm-thick Al mask (by lift-off using AZ 5412E, and etching by O_2_ plasma. Afterwards, the graphene grown by chemical vapour deposition on Cu was transferred (process done by Graphenea). Graphene was patterned by oxygen plasma (50 sccm, 300W for 1min) in a reactive ion etching (RIE). The photodefinable resist used to protect the graphene in the channel region was HIPR 6512. After the graphene etching, a second metal layer was patterned on the contacts following the same procedure as for the first layer, but this time performing an ozone cleaning treatment before metal deposition in order to improve the contact resistance and noise^133^. Subsequently, the transistors were insulated with a 3-µm-thick photodefinable SU-8 epoxy photoresist (SU-8 2005 Microchem), keeping uncovered the active area of the transistors channel. SU-8 is known to limit the stability of devices in physiological conditions to a semi-chronic time scale^134,135^. Finally, the polyimide substrate was structured in a reactive ion etching process using a thick AZ9260 positive photoresist (Clariant) layer as an etching mask. The neural probes were then peeled off from the wafer and placed in a zero insertion force (ZIF) connector to be interfaced with our custom electronic instrumentation.

### Recording systems design

We used two recording systems based on two different design approaches; the first is based on an application specific integrated circuit (ASIC)^43^ and the second on commercially available discrete components^45^. The ASIC-based approach enables the integration of the system in a fully implantable headstage and optimizes the operation for higher channel count graphene probes, while the system based on discrete-components relaxes spatial constraints on the design and facilitates early adoption of the technology. To cope with the large dynamic range of the DC-coupled signals, including the variability in the DC current offsets between graphene transistors, both design approaches present a low gain (10x) transimpedance amplifier as the first amplification. To reduce the dynamic range of the signal, the ASIC-based system includes current offset cancellation circuitry with a programmable resistor array to subtract current at the transimpedance amplifier input corresponding to specific g-SGFET offsets (Supplemental Data Fig. 1). The amplifier voltage noise at the output increases with the number of columns (Supplemental Data Fig. 2). In order to minimize the contribution of *e*_*amp*_, the ASIC implements a correlated double sample (CDS), which samples the noise from the amplifier and subtracts it from the sampled signal. On the other hand, the system based on discrete components copes with the large dynamic range of the signal by digitizing it at two stages of amplification (Fig. S1b-left). The output from the first stage is digitised after the low gain transimpedance amplifier as the DC-coupled signal. The analog signal after the first stage is high-pass filtered above 0.1Hz to eliminate common offsets and further amplified before digitization resulting in the effective prevention of quantization noise.

After amplification, the ASIC implements the analog-to-digital conversion (ADC) and the serialization for effective data transmission. The TDM read-out integrated circuits (ROIC) have been combined with an electrophysiology recording platform from Multi Channel Systems (MCS). The setup is based on an interface board (IFB) and a signal collector unit (SCU) (Supplemental Data Fig. 1). The IFB provides a USB 3.0 interface with the computer, while the SCU provides a bridge between the field-programmable gate array (FPGA) on the headstage (built around the LFE5U-45F device from Lattice Semiconductor) and the IFB itself. On the other side, a custom-made cable ends in an 8-pin polarised connector PZN-08-AA from Omnetics connector and carries the 4 differential wire pairs needed for the master-slave data input and output, clock and supply to the headstage. The headstage is composed of two PCB modules, the ROIC carrier and the FPGA board, which can manage several ROIC modules. The FPGA already contains several IPs and all the necessary blocks for the ROIC application, such as the 51.2 MHz phase-locked loop (PLL), a mapped block random access memory (BRAM) module, the high-frequency serializer/deserializer (SerDes) and the low-voltage differential signalling (LVDS) drivers for the system cable. The complete hardware setup is configured with the help of the open-source Scientific Python environment and a proprietary dynamic-link library (DLL) from MCS. The system based on discrete components is based on the conduction of the analog signals (as changes in the drain-to-source current) via 32 wires to a NI-DAQ card for digitisation and communication with the PC.

### Ethical approval and animal handling

The experiments in-vivo were in accordance with the European Union guidelines on the protection of vertebrates used for experimentation (Directive 2010/63/EU of the European Parliament and of the Council of 22 September 2010). Electrophysiological experiments with Long Evans rats were carried out under the German Law for Protection of Animals (TierSchG) and were approved by the local authorities (ROB-55.2-2532.Vet_02-22-22). Rats were kept under standard conditions (room temperature 22 ± 2 °C, 12:12 h light–dark cycle, lights on at 10:00), with food and water available ad libitum.

### Surgical implantation of large-scale graphene ECoGs

A cohort of 8 Long Evans rats, weighing between 558±121 g were used for this study. As previously described^42^, anaesthesia was induced with Midazolam (2mg/kg), Medetomidin (0.15mg/kg) and Fentanyl (0.005mg/kg). After 1h of surgery, isoflurane at a concentration of 0.5-1.5% was provided while monitoring vital signs. The posterior-dorsal area of the head was shaved, the skin was locally disinfected with Povidone-iodine and subcutaneously infiltrated with the local anaesthetic Bupivacaine. The skin was then incised and the dorsal skull was cleaned carefully by blunt dissection and the temporal muscle was partially separated from the skull. The skull was cleaned with H_2_O_2_ (3%) and phosphoric acid (KERR - Gel Etchant, 37% phosphoric acid), sequentially for a few seconds and then by scratching the surface with a scalpel. A 3D printed base ring made of Polylactic acid (tough PLA, Ultimaker) was anchored to the skull with at least six screws in the frontal side of the skull, two more on each temporal plate of the skull, two on the occipital plate and with Metabond cement (Parkell).

After at least 7 days of recovery, the animals underwent a second surgery for the implantation of the neural probes. This two-step surgery ensured fast recovery following the probes implantation. The animals underwent the same anaesthetization procedure described above, with Metamizol (110mg/kg) provided to maintain the analgesic effect 4h after the start of the surgery. A small craniotomy (∼1mm x 2mm) was opened above the cerebellum for the placement of an Ag/AgCl reference electrode (100µm thick) close to the cerebellar surface. In order to minimize the contact of Ag with the tissue and improve the electrochemical stability of the reference, the electrodes and the cerebellar surface were covered with a drop of agar solution (0.5%). Secondly, a large craniotomy was opened on the frame defined by the base-ring with a width of 1.1cm and length of 1.3cm. To perform such a large craniotomy, the skull was removed in five pieces. Four symmetric quadrants and the skull on the midline. For the detachment of the skull from the dura, a blunt tool was used, with special care on the area around the sinus. Afterwards, the dura was removed within the area of the craniotomy and the device was placed in its final position. The craniotomies were covered with a pre-polymerised PDMS (∼100µm thick – SYLGARD 184) and a 3D-printed skull replacement was lowered using the stereotaxic micromanipulator until it was in contact with the PDMS and then lowered by ∼200µm to correct for the mild brain bulging. The gaps around the skull replacement were filled with fresh Dura-Gel (Cambridge Neurotech) to protect the brain surface and keep the depth probe in place. After 15 minutes for Dura-Gel curation, the craniotomy was completely sealed using UV-curable cyanoacrylate-based glue (Loctite 4305).

### Implantation of flexible depth probes

For the insertion of flexible depth probes, rigid silicon shuttles with polyethylene glycol (PEG) 10000MW were used as previously reported^48,49^. Flexible depth probes were aligned with the silicon shuttles with previously added PEG using a micromanipulator under the microscope. When properly aligned, the PEG was molten on a hot plate. To have finer control of the placement of the connectors during implantation, the shuttle was attached to an additional motorised stage on the stereotaxic frame. After insertion of the depth probe into its final location, the PEG was dissolved using physiological saline solution. Once the flexible probe was delaminated from the shuttle, it was immobilised using a thin layer of agar (0.5% mass) after cooling down (shortly before solidifying). Subsequently, the additional motorised stage was used to extract the shuttle without moving the flexible probe or its connector. Finally, the probe was fully immobilised using a layer of Dura-Gel (Cambridge Neurotech) to protect the brain followed by encapsulation using optical cement (Evoflow - Ivoclar Vivadent, Liechtenstein) and the connector moved to its final position using the stereotaxic frame.

### Electrophysiological recordings

Electrophysiological recordings using the large-scale graphene probes and depth graphene probes were carried out in a circular open field arena (0.8m in diameter) with high walls transparent to IR light for motion capture tracking. The recordings lasted between 46 minutes and 248 minutes (mean of 145.8 minutes). The discrete electronics system was based on a previous version^45^, upscaled to enable the operation of up to 256 channels. Shortly, the system consists of a commercial data acquisition card (National Instruments DAQ-Card, USB-6363), and a custom-built amplification front-end and switching array for multiplexing (see Supplemental Data Fig. 1). The sampling frequency of the DAQ-Card of 1MHz is shared among the 256 channels, the DC-coupled and AC-coupled amplification stages. For each switch between columns 2 samples are dropped for signal stabilization leading to a sampling frequency of 651Hz each channel. This system was operated using the code published in the repository (https://github.com/RamonCortadella/LargeScale-WideBand/) for the acquisition of calibration *I*_*ds*_ − *V*_*gs*_ curves as well as for brain mapping. In the second system, an ASIC^43^In the second system, an ASIC^43^ implements the switching matrix, I-V conversion with offset cancellation and further amplification and digitization with a correlated double sampling. An FPGA on the headstage implements the digital communication between the ASIC and an interface board (IFB, Multi-Channel Systems and MCS-SCU-in-vitro, Multi Channel Systems) to implement the data transmission to the headstage. The system was operated using customised software from Multi-Channel Systems. The sampling frequency was 12 kHz per channel, way above the LFP band, to reduce high-frequency noise (see Supplemental Data Fig. 2).

### Signal pre-processing

The acquired signals representing the drain-source currents through all transistors in the active matrix were calibrated using the stationary transfer curve (*I*_*ds*_ − *V*_*gs*_) by interpolating the DC-coupled current. Using this method, variations in the *G*_*m*_ due to *V*_*gs*_ drifts are corrected^136^. Following interpolation, the signals were high-pass filtered using a median filter with a time constant of 200s, which preserves better changes in the abrupt changes in the DC-potential level compared to IIR filters. The signals were converted to a binary format (16-bit) with a range of 10mV for full-band recordings (ASIC-based system), 30mV for DC-coupled channels (discrete-components-based system) and 3mV for the AC-coupled channels (discrete-components-based system). Bad channels were then detected as outliers (x330 or /330 times the power) in the low frequency (1-10Hz) or high frequency (60-120Hz) within neighbouring 3×3 pixels. The bad channels were then interpolated using the “scatterinterpolant” function in Matlab, and was median-filtered with a 3×3 kernel and using constant padding.

### Data analysis

#### Spectrogram calculation

Spectrograms were computed using the multitaper method on the whitened signal. Whitening was performed using a first order autoregressive model^137^Whitening was performed using a first order autoregressive model^137^ fitted on the LFP in the 1-100Hz band (excluding the 50Hz and 100Hz pick-up noise peaks by a notch filter). The window length for the FFT was 2.36s (or 2.46s for the 512 channel probes). The overlap between FFT windows was half of the FFT window duration and time-frequency bandwidth NW was 3, corresponding to 1.27Hz frequency bandwidth. The spectrograms for factor analysis of sleep spindle power were computed with a window of 1.576s, 7/8 window overlap, NW=2, frequency bandwidth of 1.27 Hz.

#### Brain states classification

Brain state classification was based on the combination of behavioural states obtained from motion tracking and electrophysiological states obtained from the high channel-count neural probes (see Results section). Motor state classification was based on thresholding the speed magnitude (with a speed of 5mm/s) to define periods of activity and inactivity (>5s) and motor-defined “sleep” starting after 20s of inactivity and lasting at least 20s more). The time series of transitions between the electrophysiological states (i.e. SWS, theta and HVS) classified from spectral characteristics were smoothed using a median filter to eliminate exceedingly fast transitions. SWS and theta states were filtered with a time constant of 20 s and HVS states with a time constant of 1.5s. Then, the electrophysiological states were combined with the motor state for complete brain state classification. SWS was defined as overlapping periods of sleep and SW activity. Awake non-theta (Aw. NTHE) states were non-sleep periods overlapping with SWS states. THE (exploratory state) consists of overlapping theta and active state. For REM sleep classification, the active/inactive state transition time series was median filtered with a time constant of 20s to exclude brief movements during REM and preserve the continuity of the state. Electrophysiological states classified as theta that were overlapping with presumed sleep states were classified as REM when preceded by SWS. HVS events were defined solely based on the electrophysiological classification. Finally, MA were defined as bouts of active state shorter than 10s within 40s of inactivity and limited to SWS (with these short bouts of movement not interrupting the SWS epochs).

#### Measurement of separation between state clusters

Quantification of the separability of brain states was evaluated using rescaled Mahalanobis distances. More precisely, the Mahalanobis distances for all data points with respect to the points in a particular state were computed. Then, the Mahalanobis distances of all points that do not belong to that state were rescaled and summed. The rescaling was performed according to the complementary of the cumulative chi-squared distribution with 2 degrees of freedom (number of dimensions in UMAP embedding). Using this rescaling, those points that are nearer to the cluster of the particular state, but do not belong to it, have a higher contribution. The summed distance divided by N is the *L-ratio* used to quantify the separability of clusters resulting from the automatic spike-sorting, a procedure analogous to state-sorting implemented here^138^. The larger the *L-ratio*, the less isolated is the given cluster.

#### Computation of mean spectrum across cortical regions and states

The mean spectrum was computed from the spectrogram across samples for each brain state to provide spatio-spectral characteristics of each state. The mean spectrum was re-whitened in each state separately by performing a linear detrend on the log-transformed mean spectrum. The linear fit from a single reference channel (second row and second column) was used for all channels. The log-transformation was reversed by raising the values to the exponent (the result is expressed in arbitrary units). To compare the power across animals, we computed the power index as the median of the band-integrated power across ML axis, subtracted the mean across AP axis and normalised by its range (maximum to minimum). The mean spectrum during immobility in the infra-slow band was computed including wake immobility and sleep as well as short periods (<100s) of locomotion in between.

#### Independent component analysis of wide-band LFP around HVS

LFP was band-pass filtered in the infra-slow part via low-pass filtered below 0.5Hz and the oscillatory part capturing power in of the spike and wave activity by filtering in the 5-30Hz band. Epochs of multichannel time series for these two bands centered around the onset of HVS (±20s) were detrended, their median subtracted, concatenated and ICA (FastICA^139^) was applied to the resulting time series giving rise to two sets of independent components (ICs), IC_lp_ and IC_bp_. The activation dynamics of all ICs were used to reconstruct the contribution to the LFP from each component by multiplying the maximum loading value and its activation score. To sort the ICs, the mean reconstructed LFP (and RMS) at the site of largest loading was computed for the 2s time window preceding and following the HVS onset. The DC-potential shift and RMS was used for sorting the ICs in the infra-slow and spike and wave activity bands respectively. The loadings of the first component were compared across animals after normalizing the loadings by their value at the channel located - 3.5mm with respect to bregma (AP axis) and 3.5 with respect to the midline (ML axis).

#### SVD of infra-slow LPF and analysis of temporal relation to state transitions

The LFP during SWS and REM plus the preceding and following the 20s was concatenated across sessions for each animal, filtered below 0.5Hz and rank transformed to normalize and reduce the impact of motion artefacts on the decomposition. The SVD of the transformed signal revealed the common second component (SV2) across animals with consistent anteroposterior and mediolateral gradient. A consistency of the 2^nd^ SV component loadings across all animals was quantified by summing up the sign of the left-singular vectors across all animals, which would give +/−6 if the loading has a consistent sign across all animals (see Fig.4b bottom). Infra-slow events characterized by this component were detected as troughs in the SV2 score timeseries with the prominence above 90th percentile. These events were used to compute the inter-even intervals and cross-correlation with the MA.

#### Triggered average of isLFP and spindle power around states onset

To compute the triggered average, the LFP was low-pass filtered below 0.5Hz and the signal within the MA (0.7s post onset) or REM sleep (10s post-onset) was subtracted in the time window of interest (±30s for MA and ±150s for SWS/REM onset).

#### Identification of putative down-states for depth probe position estimation

The LFP signal during SWS from ECoG recordings was filtered in the delta band (0.5-3Hz) and the mean across all channels was used to detect global delta waves. Delta waves were detected as large amplitude troughs in the mean surface LFP exceeding 2 standard deviations. This procedure was validated using concurrent depth recordings where delta waves consistently presented a phase reversal across layers (see Fig. 5h and Supplemental Data Fig. 11), in accordance with prior literature^73^.

#### Factor analysis for topographic sleep spindle detection

Factor analysis was performed as previously described^52^. Briefly, it consists of the following steps. First, the power per frequency bin and channel is log-transformed to bring the distributions closer to symmetric (normal) and then z-scored to have homogeneous variance across all dimensions. The spectrogram is also spatially smoothed in each frequency bin and time point using a 3×3 median filter across pixels. Then principal component analysis is performed and 99.9% of the variance is kept. Factor orthogonal rotation was then performed using the Varimax method^140^, using the simplicity criteria to obtain a matrix of factor loadings. The simplicity criterion finds directions with the most parsimonious structure in the covariance matrix, which have high loadings in only a few spatio-spectral variables and close to zero in most of them. In addition, factor loadings are all positive which eliminates the ambiguity in the polarity of principal components.

#### Specificity calculation

The spectrogram in the sigma band was spatially smoothed (using a 3×3 median filter across pixels) for each frequency bin and time point. Then the spatial-frequency coordinates with a loading above half of the maximum were taken as the signal and those below as the noise. The ratio between the mean power of the signal and the mean power of noise was defined as the specificity of a factor to describe the covariance structure of the spectrogram in the sigma band.

#### Sleep spindle duration estimation

The duration of sleep spindles was computed from the activation dynamics of distinct factors. The activation, analogous to ICA activations or PCA scores, reflects the projection of the log-transformed, z-scored spectrogram on the direction of the loadings. This projection does not reflect a measure of reconstructed power but of log-transformed, z-scored power. The log transformation was reversed, effectively stretching positive deviations, and the duration was defined as the full width at half maximum (FWHM) of the activation peaks. The traces shown in Fig. 5b correspond to the activations after reversing the log transformation.

#### Matching of factor loadings

For comparison of factor characteristics across animals, factor loadings from different animals are matched based on their Euclidian distance in a 7-dimensional feature space. The features used for matching are the sign in the ML axis, the sign in the AP axis with respect to the centre of the array, the actual position in the array in units of pixels (X and Y axis), maximum frequency in rescaled frequency axis (centred at 0 and normalised by the maximum variation of 5Hz). The number of channels with loadings above half of the maximum loading and the number of frequency bins with maximum loading above half of the maximum (both normalised by the number of channels in a quadrant of the grid) were also used.

#### Harmonic index calculation

The spectrogram was log-transformed and z-scored to be in the same space used for factor rotation. Then its value was taken at all sleep spindle occurrence times (i.e. times of peaks in the activation of individual factors) at the location and at the frequency of peak of the respective factor loadings. The power at the peak frequency minus the mean of the ±5Hz points defines our measure of harmonicity. The harmonic index can be interpreted as the number of standard deviations between the power at the frequency of maximum loading and the power at 5Hz below and above (in the z-scored and log-scaled power space).

#### Temporal coupling between infra-slow networks activation and sleep spindles

The time difference between each of the spindle events from all factors and the troughs in the isLFP in an AL region of interest (−4.2mm, −1.5mm / ML, AP) were computed. Then, the JPDF for lags and the location of the spindle oscillator (defined as the quadrant with maximum loading using polar coordinates centred on the midline and at - 4mm from Bregma in the AP axis) was computed. The same distribution was computed from randomised time points within SWS periods for the gradient peaks and the JPDF recomputed from 10 randomizations. The mean JPDF from the randomised distributions was subtracted from the non-randomised JPDF in order to eliminate the edge bias produced by having gradient peaks close to the onset or offset of SWS epochs.

#### Coupling between slow oscillations and local sleep spindles

To evaluate the coupling between SO and sleep spindles we first evaluated the phase statistics of global SO at the time of peaks in scores of the first 10 factors for all animals by taking the phase of the mean LFP signal across the entire array. To show the preferred phase and strength of modulation for all animals separately we computed the resultant vector for the first 10 factors independently on each of the 8 animals. Once the temporal coupling was established, we looked into topographical coupling between SO and sleep spindles by calculating the correlation between the amplitude of the SO in all channels of the array and the amplitude of sleep spindles for each of the first 15 factors and each of the 8 animals. The amplitude of the slow oscillation was determined by finding the preceding trough (between 0.8 and 0.2s of the spindle time) and the peak (within 200ms of the spindle time) at the location of maximum loading. The SO amplitude at these times was computed for all channels and their correlation to the spindle score amplitude was calculated.

#### Statistical methods

Mean or medians are given with standard deviation. Significance testing of deviations from zero was done by the Wilcoxon signed-rank test. The statistical significance of the positive and negative deviations from zero observed in the triggered average was assessed by a permutation test for each pixel and time point. The sigma power median across events was compared by the Wilcoxon rank-sum test to the shuffled distribution across all frames and pixels. To take into account the false discovery for such a large number of comparisons we applied the Benjamin and Hochberg correction both for the LFP deviations from zero and sigma power deviations from the shuffled distribution^141^. To evaluate the topographical coupling between sleep spindles and SO the correlation maps and the factor loadings were compared by computing the correlation between these two maps in the hemisphere of maximum loading. To check for the significance of the correlation, the pixels in the correlation map were shuffled and a p-value for the correlation was obtained (1000 permutations).

## Data availability

The characterization of devices in-vitro and the complete electrophysiological and motor tracking dataset is available from the corresponding author upon request. Movies can be found in (https://doi.org/10.12751/g-node.kv2mlt).

## Code availability

The complete code for analysis of electrophysiological data is available from the corresponding author upon request. Code for the operation of the custom-built recording system can be found in https://github.com/RamonCortadella/LargeScale-WideBand/

## Acknowledgments

We thank Roustem Khazipov and members of Sirota lab for useful discussions. This work has been funded by the European Union’s Horizon 2020 research and innovation program under Grant Agreement No. 732032 (BrainCom), Grant Agreement No. 881603 (GrapheneCore3). Research at the LMU was supported by the Bundesministerium für Bildung und Forschung [grant number 01GQ0440]. ICN2 is supported by the Severo Ochoa Centres of Excellence programme, funded by the Spanish Research Agency (AEI, grant no. SEV-2017–0706), and by the CERCA Programme/Generalitat de Catalunya. RGC received funding from the Alexander von Humboldt foundation and the Marie Curie individual fellowship program. This work has made use of the Spanish ICTS Network MICRONANOFABS, partially supported by MICINN and the ICTS ‘NANBIOSIS’, more specifically by the Micro-NanoTechnology Unit of the CIBER in Bioengineering, Biomaterials and Nanomedicine (CIBER-BBN) at the IMB-CNM. We also acknowledge funding from the Generalitat de Catalunya (2017 SGR 1426), and the 2DTecBio project (FIS2017-85787-R) funded by the Ministerio de Ciencia, Innovación y Universidades of Spain, the Agencia Estatal de Investigación (AEI) and the Fondo Europeo de Desarrollo Regional (FEDER/UE). Part of this work was co-funded by the European Regional Development Funds (ERDF) allocated to the Programa operatiu FEDER de Catalunya 2014–2020, with the support of the Secretaria d’Universitats i Recerca of the Departament d’Empresa i Coneixement of the Generalitat de Catalunya for emerging technology clusters devoted to the valorisation and transfer of research results (GraphCAT 001-P-001702). Ramon Garcia-Cortadella received funding from the Alexander von Humboldt postdoctoral fellowship programme and HORIZON-MSCA-2022-PF programme.

## Author contributions

RGC contributed to the conceptual development of graphene recording systems, fabrication and characterization of graphene probes, development and execution of surgical procedures, acquisition of electrophysiological recordings, data analysis and manuscript preparation. JCF designed and characterised the ASIC and developed the headstage hardware, data communication protocols and data acquisition software. GS developed the techniques for the chronic implantation of large-scale probes. AS contributed to the factor analysis of sleep spindles. AU contributed to the data pre-processing and analysis of infra-slow dynamics. NS contributed to the conceptual design of graphene recording systems and fabrication of graphene probes. JA contributed to the development of the discrete electronics system for the operation of graphene probes. EMC contributed to the conceptual development of the technology. EDC contributed to the development of the graphene flexible electronics. RM development of electronics for ASIC headstage. JP and MK implemented the firmware for digital data communication. HL developed the software for recording and analysis. CJ and JM tested and validated the operation of the ASIC headstage. XI contributed to the design and fabrication of graphene probes. FSG led the team for the design of the ASIC and headstage. AGB and JAG coordinated the development of graphene recording systems. AS contributed to the conceptual development of the technologies and experiments. Supervised the electrophysiological recordings, data analysis and manuscript preparation.

