## Supplemental data for "DC-coupled, 2.5D electrophysiological imaging of large-scale cortical dynamics"

### Supplemental data Fig. 1: Recoding systems based on multiplexed graphene active probes

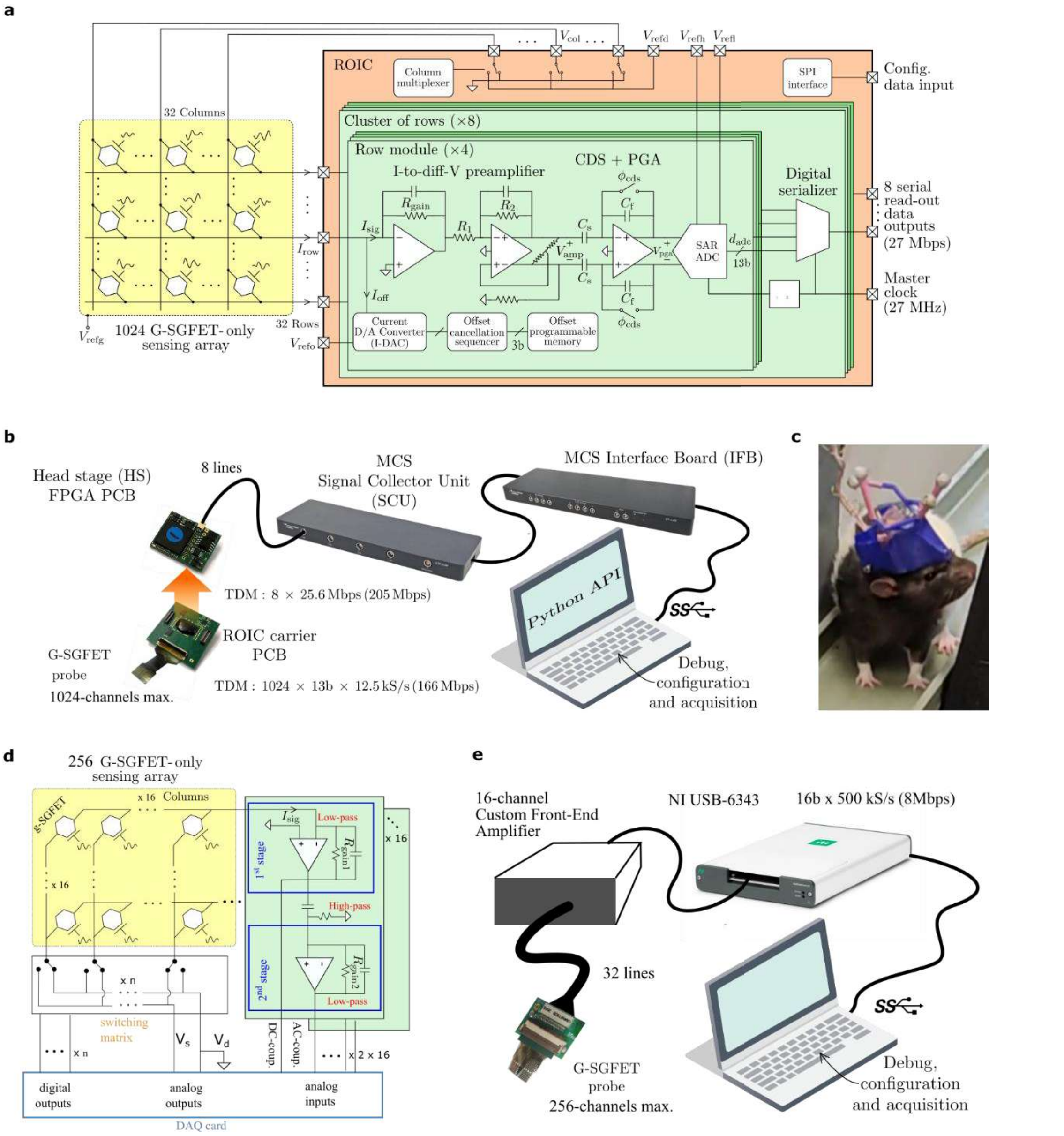


**a.** Schematic of the ASIC. **b.** Blocks diagram of the ASIC-based recording system. **c,** Photograph of a rat implanted with a 256-channel array during recordings. A reflective marker was attached for motion tracking. **d,** Schematic of recording system based on discrete electronic components. **e**. Blocks diagram for the recording system based on discrete electronic components.

**Supplemental data Fig. 2: Noise characteristics of 1024, 512 and 256 channel graphene probes operated with the recording system**

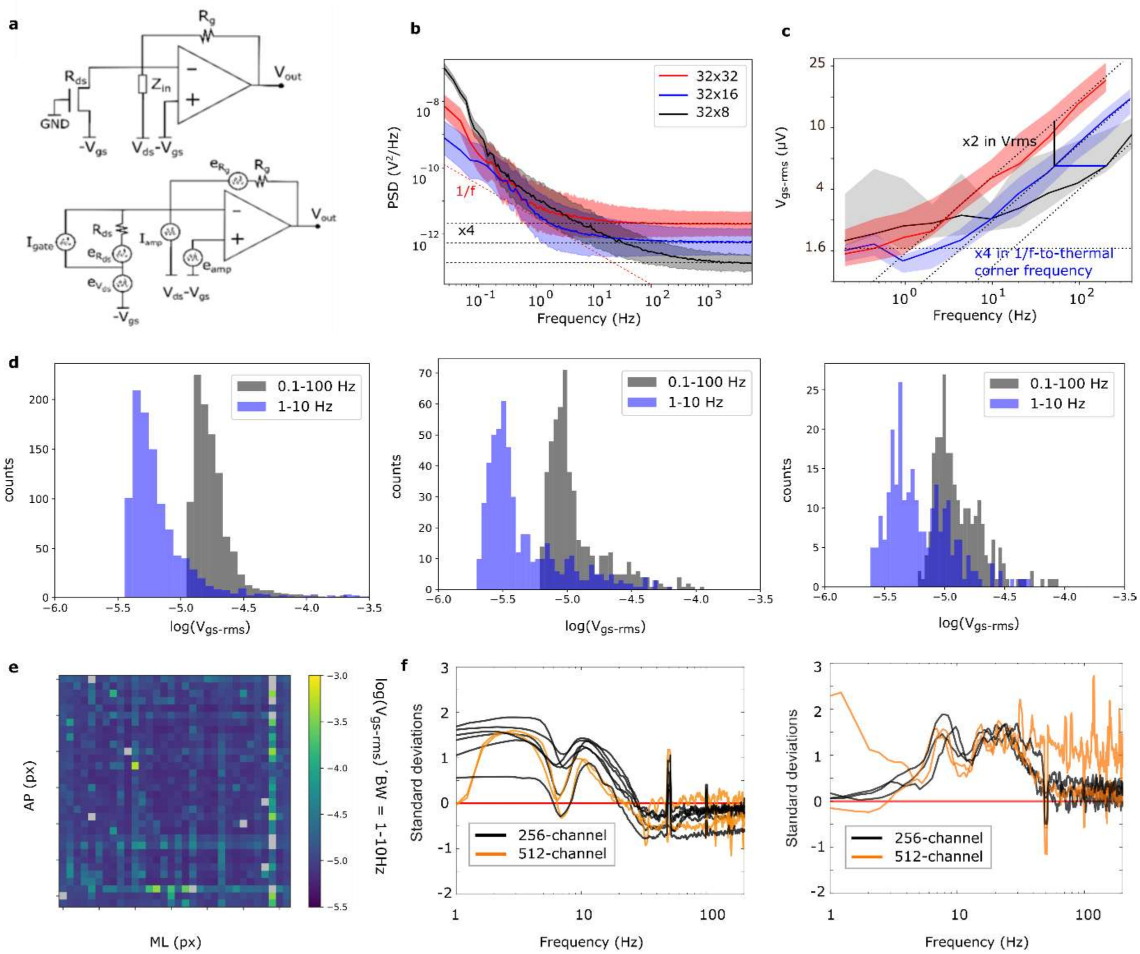


**a.** Schematic of the graphene transducers and transimpedance amplifier (top) and the noise sources in the circuit (bottom). The amplifier voltage noise ($e_{amp}$) is proportional to the amplifier gain and inversely proportional to the load impedance in the limit of small load values. **b.** Power spectral density mean (solid line) and standard deviation (shaded area) for a 32x32 transistor array, a 32x16 array and a 32x 8 array. Note the increase in floor noise due to increased aliasing and increased amplification of the intrinsic voltage noise of the amplifier. **c.** The equivalent noise at the gate (Vgs-rms) for a sliding frequency window of [fmin-fmin*$10^{0.5}$] (i.e. half the Vgs-rms in a frequency decade for 1/f noise). **d.** Vgs-rms across channels for a 32x32 transistor array (left), 32x16 transistor array (middle) and 32x8 transistor array (right) integrated in a wide band (0.1-100Hz) and in a one frequency-decade band (1-10Hz). **e.** Vgs-rms integrated in the 1-10Hz band across channels mapped onto the 2-dimensional array for a 1024 channel array **f.** The difference in the z-scored mean spectrum between SWS and THE (left) and HVS and awake non-THE (right) shows that above ~50 Hz the measured difference in power between states is relatively small compared to lower frequencies.

**Supplemental data Fig. 3: Sensitivity of graphene probes before and during chronic implantation**

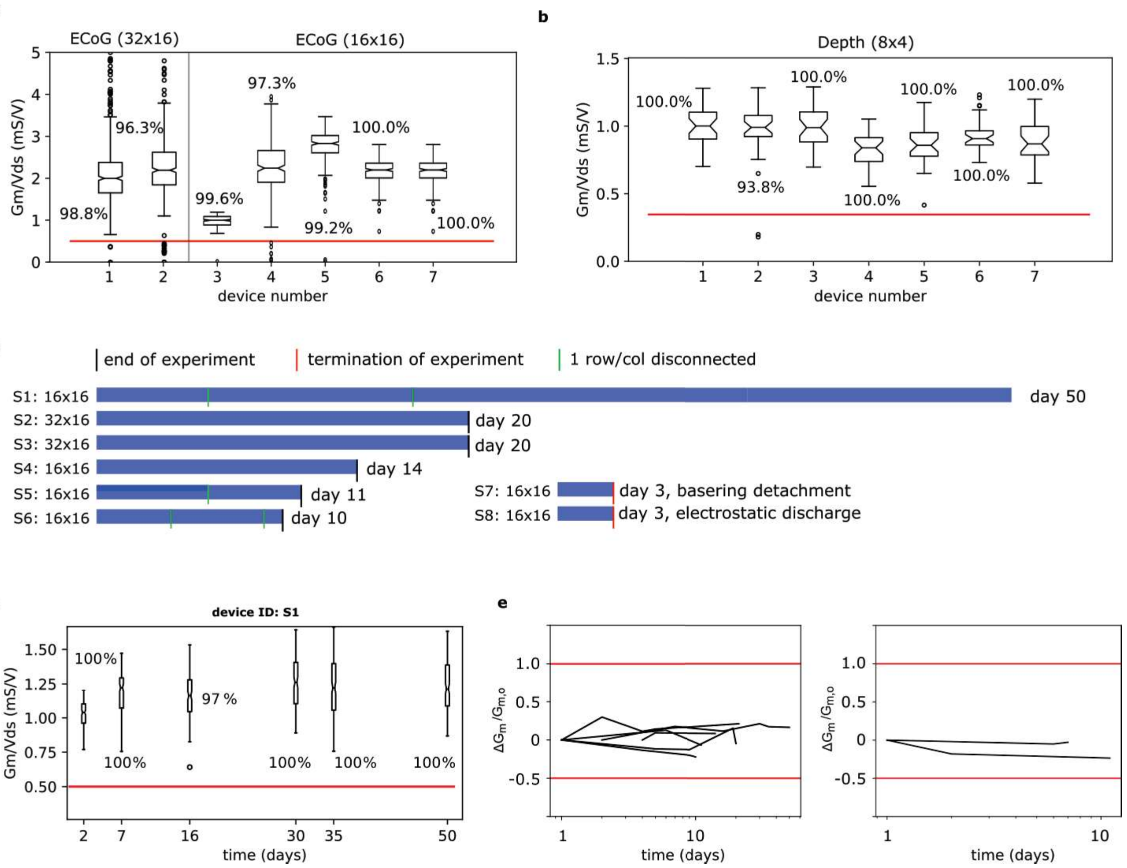


**a.** Normalised transconductance at optimal gate potential for all the ECoGs used in this study measured in saline solution**. b.** Normalised transconductance from 8 depth graphene probes, each with 32 channels. In panel a and b the yield is estimated as the number of channels with a transconductance >0.4mS/V, which is labelled for each boxplot. **c.** Summary of the devices implanted and the number of days they remained implanted. Two experiments were terminated earlier because of mechanical failure of the base-ring or because of an electrostatic discharge. The green vertical lines mark the disconnection of a column/row because of contact problems at the connector. **d.** Normalised transconductance for all transistors in the 256-channel ECoG S1. The yield over time is estimated as the number of channels above 0.5mS/V. **e.** The change in Gm from the first day normalised by the Gm on day 1 for all functional GFETs in devices S1-6 (left) and in the two depth probes (right).

**Supplemental data Fig. 4: Characterization of brain-states detected from spatio-spectral characteristics**

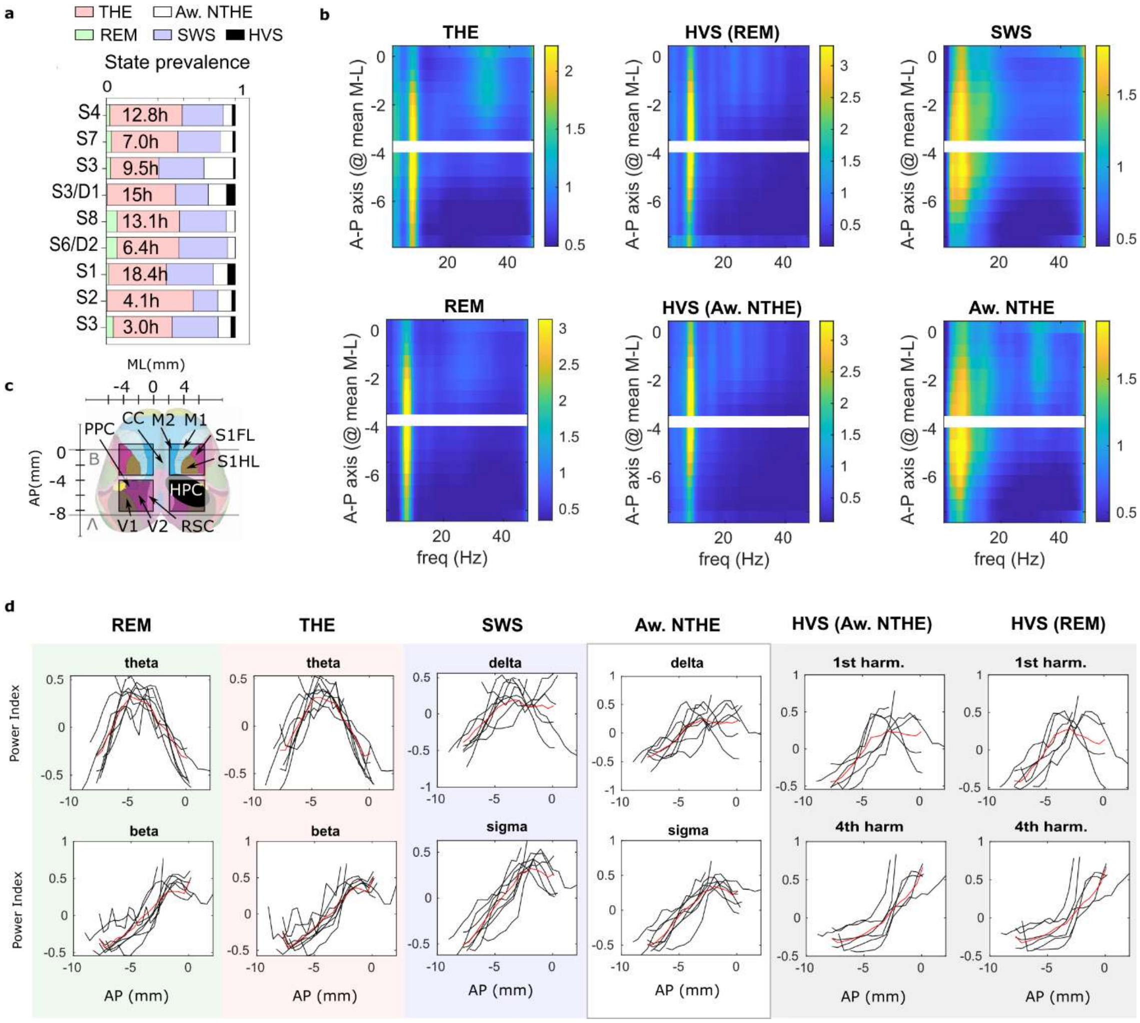


**a.** Prevalence of the detected brain states across animals. The text within the bar-plots indicates the total duration of the recordings for each animal. **b.** Mean spectrum (averaged across the medio-lateral axis, color-coded) across the anterior-posterior axis for all detected states. **b.** Mean spectrum averaged across the medio-lateral axis within various frequency bands and states (top labels). **c.** Atlas of the rat brain (obtained from EBRAINS database). Projection of the hippocampus on the surface in black. The non-shadowed area represents the sampled area by 256-channel ECoGs. ML, medio-lateral; AP, anterio-posterior. **d**. Mean spectral power (ML-axis-averaged) in the respective frequency bands (theta, 6-9 Hz, delta, 2-5 Hz; 1$^{st}$ harmonic of HVS, 14-16 Hz; beta, 20-30 Hz; sigma, 10-15 Hz; 4$^{th}$ harmonic of HVS, 30-35 Hz) across ML axis for different states (columns). Gray, single animals; black line, group mean (n=8 for SWS, REM, THE, Aw. NTHE and n=6 for HVS).

**Supplemental data Fig. 5: Relation between anterio-posterior isLFP gradient and sub-threshold motor activity**

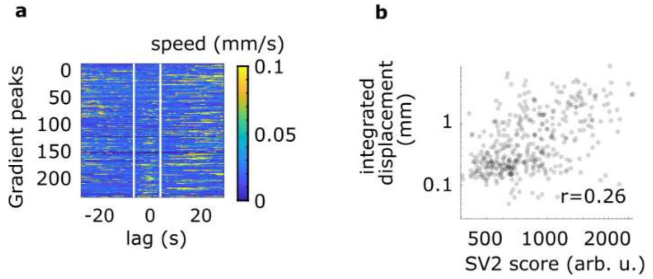


**a.** Speed aligned to the SV2 score troughs selected at least XX s away from MA to study sub-threshold micromovements associated with the isLFP dynamics. **b**. Scatter plot showing the rank correlation between the speed integrated in the (-8s to 2s) window prior to SV2 score troughs and the amplitude of the SV2 score troughs (rank correlation r=0.26, p-value < 0.0001, n=6 animals).

**Supplemental data Fig. 6: infra-slow LFP and sigma band power variations significance around micro-arousals and SWS onset**

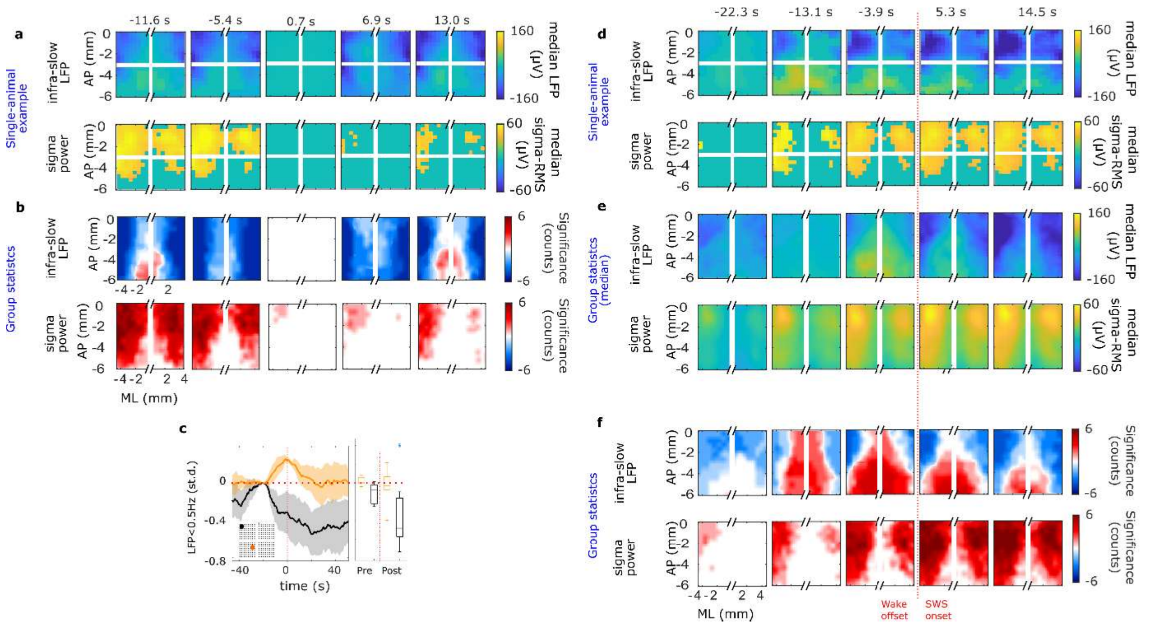


**a.** Single animal example triggered average of the LFP <0.5Hz (top) and sigma power (bottom) where non-significant pixels/frames are masked to 0. For the LFP, Wilcoxon signed-rank test ($p<0.05$, n= 6 animals) is used while for sigma power Wilcoxon Rank-Sum test between the distribution across events and the shuffled distribution across frames and pixels ($p<0.05$, n= 6 animals), both are FDR-corrected (see Methods). **b.** Sum of significant pixels across animals (n=6) for infra-slow LFP (top) and spindle RMS (bottom) around micro-arousals. **c.** Average LFP <0.5Hz, normalized by the LFP variance across the array, around SWS sleep onset (solid line is the mean and shaded area the standard deviation across animals). The boxplot shows the distribution of normalized LFP across animals for the two positions 30s before and after the SWS onset (Wilcoxon Signed-rank test, $p<0.05$, n= 6 animals). **d.** Same as panel a around the SWS onset. **e.** Sigma band RMS normalized by the RMS variance across the array around SWS onset for two example channels depicted in the inset (solid line is the median and the shaded area the standard deviation across animals, n=6 animals). **e.** Median infra-slow LFP (top) and sigma-band RMS (bottom) across animals (n=6 animals) around the SWS onset. **f.** Same as panel b around SWS onset.

### Supplemental data Fig. 7: Factor analysis of broad sigma band

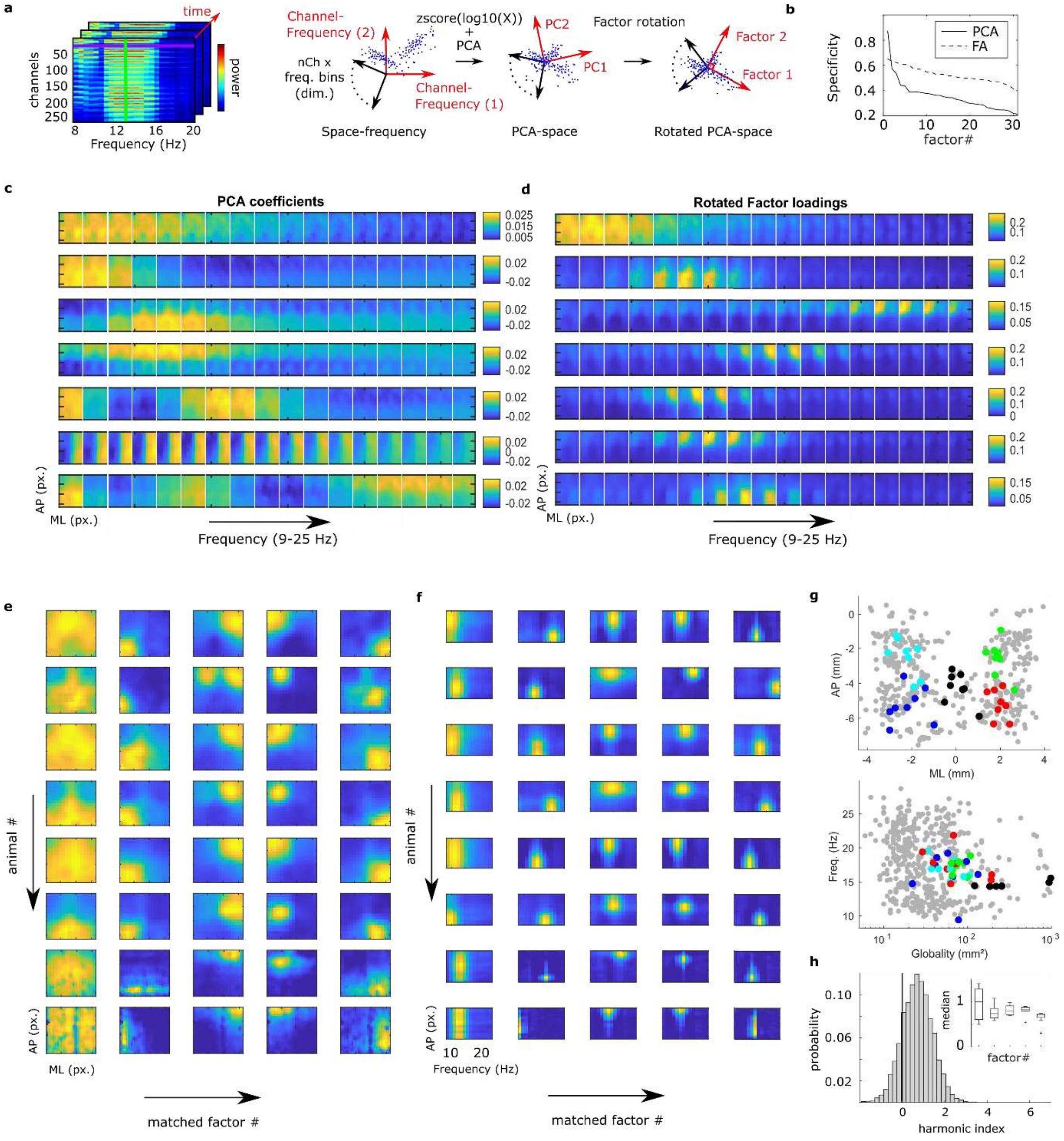


**a.** Schematic representation of spatio-spectral factor analysis. From a high-dimensional spatio-spectral space (left), factor analysis finds the most parsimonious modes of spatio-spectral covariation (right). **b.** 95 percentile of the specificity distribution across factors for PCA and rotated factors (FA) sorted by specificity. **c.** PCA coefficients in the space-frequency dimensions. Note that each rectangular color plot separated by a white vertical line corresponds to the loadings in space for a particular frequency bin. Each row corresponds to a component, from the first PC with the largest variance to the fifth component. **d.** Same representation as in c, but for the factor loadings after rotation of PCs. Note that there is no correspondence between the indices of PC in c and factors in d. **e.** Loadings in space for matched factors. Each column corresponds to one factor and each row to one animal. **f.** Loadings in frequency (x-axis) and AP axis (y-axis) for matched factors. Each column corresponds to one factor and each row to one animal. **g.** Position in space of the maximum loading location (top, same as in Fig. 5 of the main text) and corresponding representation of the peak loading of factors in the frequency space and in terms of globality (measured as the number of space-frequency bins that present values above half of the maximum). Colored dots correspond to matched factors. Colors of matched factors in g match those in Fig. 5 in the main text. **h. .** Harmonic index distribution across factors for an example animal. The inset shows the median of the harmonic index for each of the matched factors across animals (n=8 animals).

### Supplemental data Fig. 8: Topography and laminar profile of isLFP during SWS

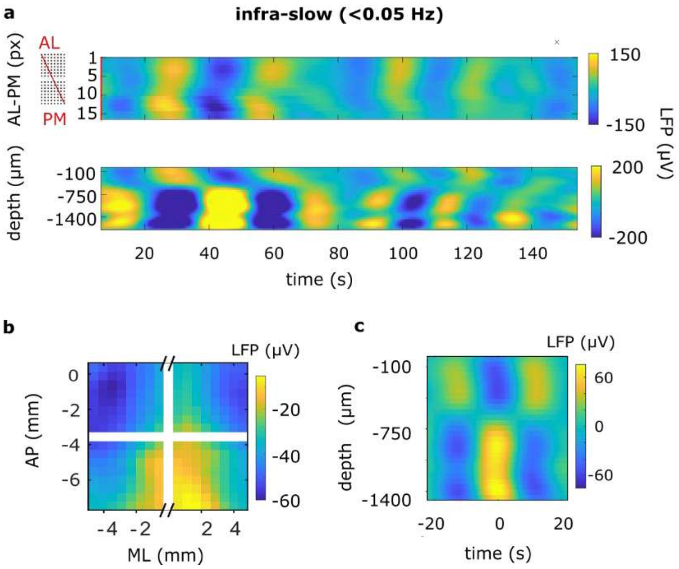


**a.** Example of the isLFP in the AL to PM axis (top) and simultaneous recording across cortical layers (bottom). **b.** Mean isLFP (<0.05 Hz) triggered at the trough in AL channels (ML -4.2mm, AP -1.5mm). **c.** Mean isLFP (<0.05 Hz) profile across cortical layers.

### Supplemental data Fig. 9: Cross-correlation between sleep spindle rate from various generators

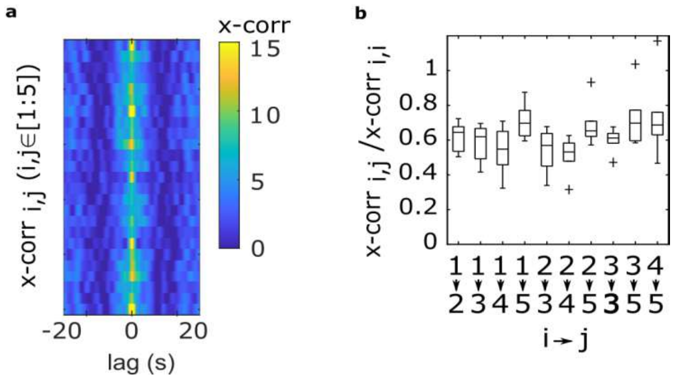


**a.** Cross correlation of the spindle events between all possible combinations of the five matched factors for an example animal. **b,** Summary of cross-correlation at 0-lag for all combinations of matched factors across animals normalised by the autocorrelation at 0-lag of the $i^{th}$ factor.

**Supplemental data Fig. 10: Correlation between preceding trough, concurrent peak, and following trough of SO to local spindles amplitude**

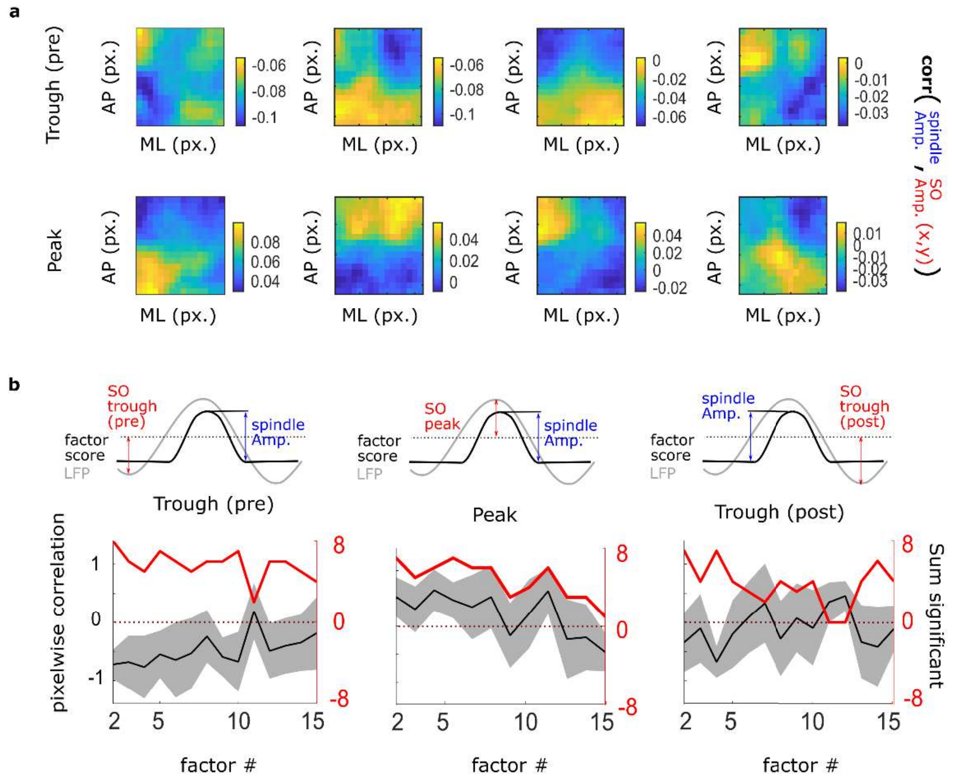


**a.** Correlation between the factor scores amplitude and the amplitude of the preceding trough (top) and concurrent peak (bottom) across all pixels of the ECoG for an example animal. Columns correspond to factors 2-4 (from left to right), matching the factors of the example animal shown in Fig. 5 of the main text. **b.** Schematic of SO trough (pre/post) and SO peak used to compute the correlation between factor score amplitude and SO amplitude at each pixel. Pixel-wise correlation (left axis, black) and sum across animals of significantly coupled factors (right axis, red) for factors 2-15 sorted by variance. Correlations for the preceding trough (left), concurrent peak (middle) and following trough (right). Significance is computed from the permutation of pixels of the correlation matrix, 1000 permutations, p-value <0.05.

**Supplemental data Fig. 11: Delta waves translaminar profile**

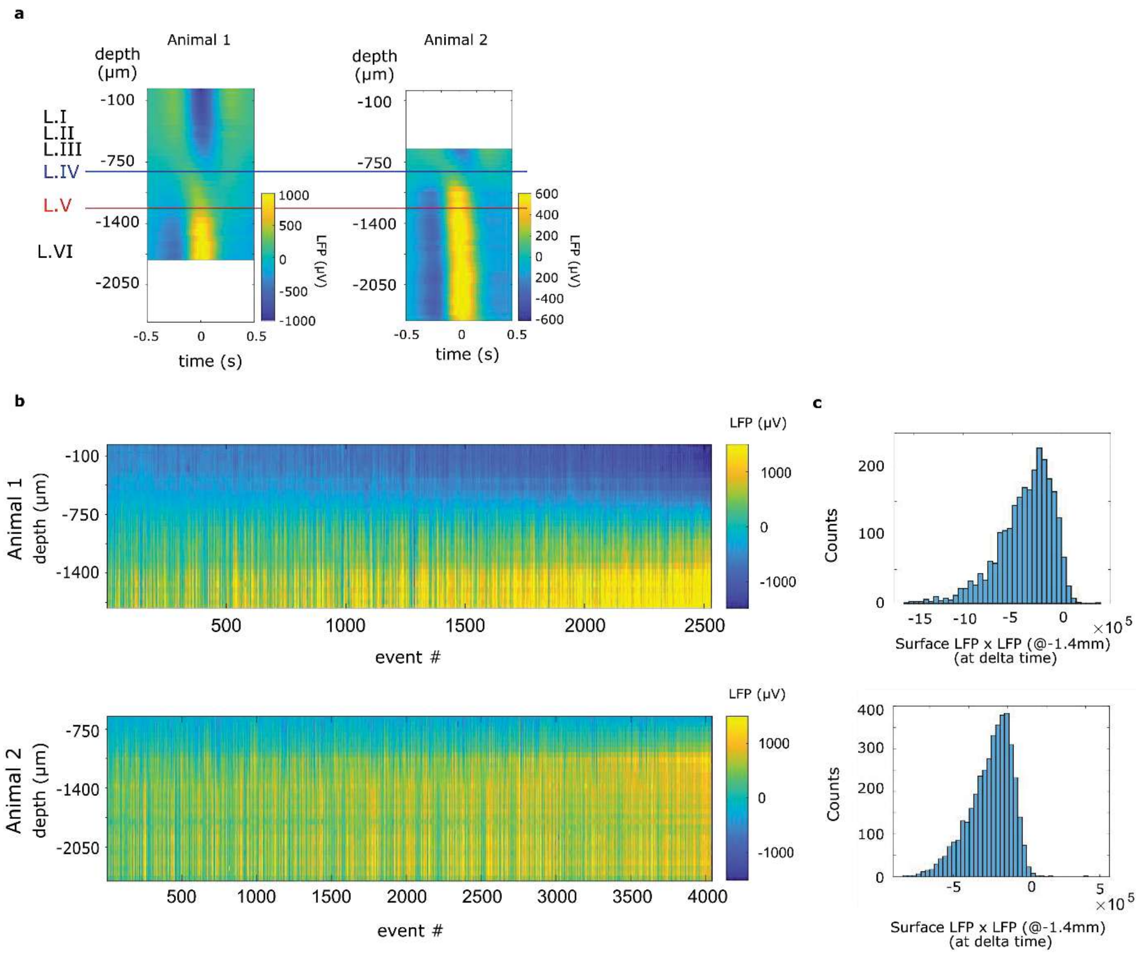


**a,** Translaminar profile of the LFP amplitude triggered on the delta waves detected on the surface for animal 1 (left) and 2 (right). Note that the scale indicating the position in the dorso-ventral axis is common for "left" and "right" plots with the position of channels shifted by approximately 600um. **b,** depth profile of the LFP a the time of SO troughs detected on the surface (>2 x s.t.d.) across events for animal 1 (top) and animal 2 (bottom). **c.** distribution of the product between the LFP at the surface and the LFP at DV = -1.4mm for animal 1 (top) and animal 2 (bottom) shows the phase reversal in the vast majority instances.
